# The identity of *Argyria lacteella* (Fabricius, 1794) (Lepidoptera, Pyraloidea, Crambinae), synonyms, and related species using morphology and DNA capture in type specimens

**DOI:** 10.1101/2022.10.10.511518

**Authors:** Bernard Landry, Julia Bilat, James Hayden, M. Alma Solis, David C. Lees, Nadir Alvarez, Théo Léger, Jérémy Gauthier

## Abstract

Recent developments in museomics enable genetic information to be recovered from previously unusable collection specimens and thus to answer complex taxonomic questions. Here we apply museomics to a taxonomic problem involving several species of *Argyria* Hübner (Pyraloidea, Crambinae), with previously unrecognized morphological variation. By analysing the DNA barcode (COI-5P) in numerous specimens, we aimed to reconstruct phylogenetic relationships between species, to provide better evidence for synonymies, and to circumscribe their geographical distribution. Using an innovative DNA hybridization capture protocol, we partially recovered the DNA barcode of the lectotype of *Argyria lacteella* (Fabricius, 1794) for comparison with the 229 DNA barcode sequences of *Argyria* specimens available in the Barcode of Life Datasystems, and this firmly establishes the identity of the species. The same protocol was used for the following type specimens: the *Argyria abronalis* (Walker, 1859) holotype, thus confirming the synonymy of this name with *A. lacteella*, the holotype of *A. lusella* (Zeller, 1863), **rev. syn**., the holotype of *A. multifacta* Dyar, 1914, **syn. n**., newly synonymized with *A. lacteella*, and a specimen of *Argyria diplomochalis* Dyar, 1913, collected in 1992. A complementary sampling composed of nine specimens of *A. lacteella, A. diplomochalis, A. centrifugens* Dyar, 1914 and *A. gonogramma* Dyar, 1915, from North to South America, were integrated using classical COI amplification and Sanger sequencing. *Argyria gonogramma* Dyar, described from Bermuda, is the name to be applied to the more widespread North American species formerly identified as *A. lacteella*. Following morphological study of its holotype, *Argyria vestalis* Butler, 1878, **syn. n**. is also synonymized with *A. lacteella*. The name *A. pusillalis* Hübner, 1818, is considered a *nomen dubium* associated with *A. gonogramma*. The adult morphology is diagnosed and illustrated, and distributions are plotted for *A. lacteella, A. diplomochalis, A. centrifugens*, and *A. gonogramma* based on slightly more than 800 specimens. For the first time, DNA barcode sequences are provided for the Antillean *A. diplomochalis*. Our work highlights the efficiency of the DNA hybrid capture enrichment method to retrieve DNA barcodes from 18th and 19th century type specimens in order to solve taxonomic issues in Lepidoptera.

## Introduction

Museum collections contain an extraordinary diversity and quantity of samples collected around the world over the past centuries (Yeates et al. 2016). Collection specimens have been used to describe and classify biodiversity on the basis of morphological criteria, but until recently it has been impossible to recover genetic information from these samples because the DNA they contain is degraded and occurs in very low quantities compared to the contaminants (Card et al. 2021). It has sometimes been possible to recover short DNA barcodes at the cost of laborious multiple PCRs (Hernández-Triana et al. 2014), but recent developments in both the ability to recover DNA, improved extractions and capture approaches, and the advent of high-throughput sequencing has opened the access to the genetic information of these specimens, allowing many new studies and the emergence of museomics (Raxworthy & Smith 2021). Among the various approaches developed to recover DNA from collection specimens, hybrid enrichment methods seem to be the most efficient (Raxworthy & Smith 2021). These capture approaches can target different regions of the genome such as mitochondria (Zhang et al. 2020), exons (Bi et al. 2012), conserved regions, i.e. conserved anchored hybrid enrichment (Espeland et al., 2018) or ultraconserved elements (UCE) (Faircloth et al. 2012; Blaimer et al. 2016), and randomly distributed loci using ddRAD approach (Suchan et al. 2016; Gauthier et al. 2020; Toussaint et al. 2021). These innovative methods make it possible to integrate old samples into modern genetic studies.

The name *Tinea lacteella* Fabricius, 1794, and its synonyms, have been applied to small white moths of the genus *Argyria* Hübner collected in the New World since Fernald (1896) synonymized with it five species: *Argyria albana* (Fabricius), *Argyria pusillalis* Hübner, *Argyria lusella* (Zeller), *Argyria rufisignell*a Zeller, and *Argyria pontiella* Zeller; Dyar (1903) added *Argyria abronalis* Walker as another synonym in a North American checklist. In the subsequent decades of the 20th century, *A. rufisignella* and *A. pontiella* were removed from the list of synonyms of *A. lacteella* while *Argyria gonogramma* Dyar, 1914 was added to it as summarized by Munroe (1995). More recently, the name *A. lacteella* has been used, for example, in Moth Photographers Group (2022), BOLD (Ratnasingham & Hebert, 2007), Scholtens and Solis (2015), and Landry et al. (2020). At the inception of this study, the BOLD database contained three widely separate lineages with specimens named *Argyria lacteella* and four with specimens identified as *Argyria centrifugens* Dyar, 1914. Among the latter group, one lineage contained specimens collected in Florida, USA; morphological examination of the holotype proved that their identification was erroneous.

Thus, because morphological and DNA barcode variation was observed in *Argyria* specimens that otherwise share a similar (ca. 11 mm) wingspan and previously unrecognized external diagnostic characters, we found it necessary to try to fix the identity of *A. lacteella* and the species similar to it, and to better understand their synonymy and geographical distribution. We aimed to do that by integrating both the COI barcode data available in BOLD and the type specimens of the species as well as those pertaining to synonymized names. For this we developed a COI barcode capture approach by designing probes along the entire barcode. We extracted historical DNA from four type specimens dating back to the 18th and 19th centuries and an additional specimen collected in 1992, and recovered COI sequence from them. We also amplified the DNA barcode for nine additional samples and integrated them with all available *Argyria* sequences in the BOLD database. Combining phylogenetic inferences, species delimitation approaches based on sequence data and morphology, we propose a new classification of several *Argyria* species. This study shows that innovative methods of museomics can solve complex taxonomic questions still debated. More generally, it reconciles the modernity of innovative molecular approaches with the biological heritage that museums have been preserving for centuries.

## Material and methods

### Sources of information

The original description of *A. lacteella* and subsequent citation of the name by Fabricius (1794, 1798) were investigated, along with the original descriptions and subsequent citations of all other taxa/names treated here. The specimens examined came from the following institutions, in chronological order of acronyms:

Carnegie Museum of Natural History, Pittsburgh, USA (CMNH);

Cornell University Insect Collection, Ithaca, New York, USA (CUIC);

Florida State Collection of Arthropods, Gainesville, Florida, USA (curated with the MGCL) (FSCA);

Museum für Naturkunde, Berlin, Germany (MFNB);

McGuire Center for Lepidoptera and Biodiversity, Gainesville, Florida, USA (MGCL);

Muséum d’histoire naturelle, Geneva, Switzerland (MHNG);

Natural History Museum, London, UK (NHMUK);

National Museum of Natural History, Washington, D.C., USA (NMNH=USNM); Oxford University Museum of Natural History, Oxford, UK (OUMNH);

Essig Museum of Entomology, University of California, Berkeley, USA (UCB);

V. O. Becker collection, Camacan, Bahia, Brazil (VOB);

Zoological Museum of the University of Copenhagen, Denmark (ZMUC).

### Dissection

Specimens from which DNA was not extracted were dissected following Robinson (1976): abdomens were macerated in hot 10% aqueous KOH, cleaned, stained variously with Orange G, chlorazol black, or eosin Y, and slide-mounted in Euparal.

### Illustrations

Photographs were taken with a variety of devices in five institutions (FSCA, MHNG, NMNH, NHMUK, ZMUC), including, at the MHNG (Figs 11-14, 20, 23-26), a Leica M205 binocular scope, a Leica DFC425 camera, and the Leica imaging software. The Visionary Digital® imaging system was used at the NMNH. At the MHNG the photos were stacked using Zerene Stacker of Zerene Systems LLC and modified for better presentation using Adobe Photoshop Elements. At the FSCA, photographs were taken with a JVC digital camera KY-F75U 3-CCD with Leica Z16 Apo and Planapo 1.0x lenses, operated and stacked with Auto-Montage Pro v. 5.01.0005 (Syncroscopy, Synoptics, 2004). High-resolution genitalic photographs (Figs 17, 18, 27, 28) were taken at the MGCL with a Leica DM6B compound microscope with a Leica DMC6200 camera, and photographs were stacked and processed with Leica Application Suite X v. 3.7.0. Postprocessing was done with Adobe Photoshop Elements 11.

### Sampling

To ascertain the identity of *Argyria lacteella*, the DNA of its unique (as far as known) type specimen housed in the ZMUC was sampled from two legs. The DNA was sampled from the abdomen of the holotype of *Argyria abronalis* (Walker, 1859), recorded as a synonym of *A. lacteella* (e.g., Munroe, 1995) and deposited in the OUMNH. DNA was also sampled from one leg of the holotype of *A. lusella* (Zeller, 1863), which had been placed as a synonym of *A. lacteella* in the NHMUK, and from one leg and part of another for the holotype of *Argyria multifacta* Dyar, 1914 deposited in the NMNH. In addition, the DNA of a specimen identified (by BL) as *Argyria diplomochalis* Dyar, 1913 from the island of Anguilla, collected in 1992 and deposited in the CMNH, was also sampled in the same manner as the old holotypes just mentioned, and the DNA of two specimens of *A. diplomochalis* collected in 2021 on Saint Croix Island, US Virgin Islands, deposited in the MHNG, was sampled from one leg each using a Sanger protocol. The four specimens used here that were sequenced at the MFNB, but deposited in the MHNG, were sampled also from one leg each with a Sanger protocol. Additional specimens were studied from the CMNH, CUIC, FSCA, MGCL, NMNH, UCB, and VOB.

### DNA extraction and capture

In the MHNG, the DNA barcode sequence from specimen *Argyria “centrifugens”* DHJ02 (BOLD sample ID BIOUG27552-D08; JEH20210604A) (in reality, *Argyria lacteella*) captured at Gainesville (Florida, USA; deposited in FSCA) (29.6922°N - 82.3650°W) was used as reference for molecular work. Probes were designed using the 648 bp reference sequence via a sliding window of 108 pb with steps of 27 bp, providing an overlap of 83 bp. Using this approach, 21 probes were designed for the forward and 21 for the reverse direction. T7 promoters were added to each probe sequence. Final probe sets were ordered from Integrated DNA Technologies (IDT). The T7 reverse-complement sequence was annealed to the probe sets to allow transcription into RNA and biotinylation in a single reaction using HiScribe T7 High Yield RNA Synthesis Kit (New England Biolabs) followed by a Dnase treatment to avoid sample contamination by probe DNA during the capture, a purification using RNeasy kit (Qiagen) and Rnase inhibition using SUPERase-IN™ (Invitrogen). Concentrations of RNA probes were measured in a Qubit RNA HS assay (Thermo Fisher Scientific).

DNA extraction on historical samples were performed using PCR & DNA Cleanup Kit (Monarch). The protocol was adapted from Patzold et al. (2020) and aims to improve the recovery of small DNA fragments on the column with the addition of ethanol. In the non-destructive protocol, after a night in Monarch gDNA Tissue lysis buffer with proteinase K (2 mg/ml final concentration), the abdomen of specimen CRA01 was treated with KOH, the genitalia were separated, and both genitalia and abdomen pelt were cleaned and mounted on slide following procedures mentioned in Landry and Becker (2021); the leg of specimen CRA02 was retrieved from the buffer and returned to the NHMUK where it is preserved in a vial underneath the specimen. In the destructive protocol the tissues were crushed (Table 1). The quality and concentration of purified DNA was assessed using a Qubit dsDNA HS assay (Thermo Fisher Scientific) and/or with Fragment Analyzer. Due to their low DNA concentration (Table 1), the samples were not diluted prior to the preparation of shotgun libraries, except for sample CRA01 (abdomen), which was diluted to ∼27 ng/μL. A modified version of the protocol from Suchan et al. (2016) used in Toussaint et al. (2021) was applied for the preparation of shotgun libraries (detailed protocol in Supplementary Material). Libraries were quantified using a Qubit dsDNA HS (Thermo Fisher Scientific) and pooled in equimolar quantities based upon their respective concentrations. For each probe set, forward and reverse, hybridization capture for enrichment of shotgun libraries was performed following the protocol described in Toussaint et al. (2021). Sequencing was performed on Illumina Miseq Nano using a paired-end 150 protocol (Lausanne Genomic Technologies Facility, Switzerland).

**Table 1.** Data pertaining to specimens sampled with hyRAD and Sanger protocols at MHNG (CRA01-07), MFNB (BLDNA 065, 137, 138, 141), and MGCL (JEH20210604A-C).

| ID | Sample | Year of collect | Extraction method | Tissue | DNA conc. (ng/ul) | #_reads | #_fragments | Mean fragment size (bp) | % COI barcode |
| --- | --- | --- | --- | --- | --- | --- | --- | --- | --- |
| *COI capture* |  |  |  |  |  |  |  |  |  |
| CRA01 | OUMNH Holotype *Argyria abronalis* (Walker, 1859) | before 1859 | non-destructive | abdomen | 53.60 | 3,100,910 | 1,559,505 | 83.25 | 99.8 |
| CRA02 | NHMUK013696753 NHMUK Holotype *Argyria lusella* (Zeller, 1863) | before 1863 | non-destructive | leg | 0.40 | 8,889 | 1,583 | 81.88 | 96.3 |
| CRA03 | ZMUC Holotype *Argyria lacteella* (Fabricius, 1794) | 1784-1789 | destructive | leg | 0.50 | 3,101 | 60 | 77.00 | 51.4 |
| CRA04 | CMNH *Argyria diplomochalis* (Dyar, 1913) | 1992 | destructive | leg | 2.94 | 862,323 | 183,175 | 103.51 | 100 |
| CRA07 | USNM *Argyria multifacta* (Dyar, 1914) | 1911 | destructive | 2 legs | 0.17 | 14,538 | 10,686 | 49.40 | 93.1 |
| *PCR amplification* |  |  |  |  |  |  |  |  |  |
| CRA05 | MHNG-ENTO-91928 MHNG *Argyria diplomochalis* (Dyar, 1913) | 2021 | destructive | leg | 0.81 |  |  |  | 100 |
| CRA06 | MHNG-ENTO-91929 MHNG *Argyria diplomochalis* (Dyar, 1913) | 2021 | destructive | leg | 0.79 |  |  |  | 100 |
| BLDNA137 | MHNG-ENTO-102922 *Argyria lacteella* (Fabricius, 1794) | 2018 | destructive | leg |  |  |  |  | 100 |
| BLDNA138 | MHNG-ENTO-102923 *Argyria lacteella* (Fabricius, 1794) | 2018 | destructive | leg |  |  |  |  | 100 |
| BLDNA141 | MHNG-ENTO-97427 *Argyria centrifugens* (Dyar, 1914) | 2018 | destructive | leg |  |  |  |  | 100 |
| BLDNA065 | MHNG-ENTO-85677 *Argyria lacteella* (Fabricius, 1794) | 2004 | destructive | leg |  |  |  |  | 100 |
| JEH20210604A | FSCA UF-FLMNH-MGCL 1112885 *Argyria lacteella* (Fabricius, 1794) | 2021 | destructive | leg |  |  |  |  | 100 |
| JEH20210604C | FSCA UF-FLMNH-MGCL 1112830 *Argyria gonogramma* (Dyar, 1915) | 2020 | destructive | leg |  |  |  |  | 100 |
| JEH20210604D | FSCA UF-FLMNH-MGCL 1112845 *Argyria gonogramma* (Dyar, 1915) | 2015 | destructive | leg |  |  |  |  | 100 |

For additional samples, the DNA barcode was amplified by PCR. For CRA05 and CRA06 (Table 1), destructive DNA extraction was performed using QIAamp DNA Micro Kit (QIAGEN) and the DNA barcode was amplified by PCR using H02198 and COImod primers (Landry & Andriollo 2020) and sequenced using Sanger sequencing (Microsynth AG, Balgach, Switzerland). Samples BLDNA 65, 137, 138, and 141 (Table 1) were processed at the MfN: DNA was extracted using the Macherey-Nagel DNA extraction kit (Dürren, Germany), and molecular work followed the protocol described in Mey et al. (2021). Sequencing was done by Macrogen (The Netherlands) in both directions. Sequences were eye-checked and aligned using Phyde 0.9971 (Müller et al., 2005). The COI barcode region of samples JEH20210604A, ∼C, and ∼D was sequenced using standard barcoding primers and protocols (Hebert et al., 2004) by the FDACS-DPI Molecular Diagnostics Laboratory, in Florida, USA.

### COI locus reconstruction and phylogeny with existing data

Raw reads were cleaned using Cutadapt (Martin 2011) to remove barcodes, adapters and bases with a low quality, and quality was first checked using FastQC (Babraham Institute). Corresponding reads were first identified by BLASTn (Camacho et al. 2009) on the reference sequence and mapped using Geneious 6.0.3 Read Mapper (Kearse et al. 2012). Consensus sequences were generated keeping the most frequent bases and a minimum coverage of 3.

### Phylogenetic inferences

To investigate the phylogeny of *Argyria* species, the sequences from all the samples including the keyword “Argyria” were retrieved from the Barcode of Life Data System (BOLD) (Supplementary Table 1). Newly generated and retrieved sequences were aligned using MAFFT (Katoh et al. 2002). Models of nucleotide substitution were identified using ModelFinder (Kalyaanamoorthy et al. 2017) implemented in IQ-TREE 2.0.5 (Minh et al. 2020). Phylogenetic inferences were performed in IQ-TREE 2.0.5 (Minh et al. 2020) and branch support were estimated using 1,000 ultrafast bootstraps along with 1,000 SH-aLRT tests (Guindon et al. 2010; Hoang et al. 2018). To avoid local optima, we performed 100 independent tree searches using IQ-TREE and selected the run showing the best likelihood score.

### Species delimitation and genetic distance

Three different methods were used to investigate species delimitation: Automatic Barcode Gap Discovery (ABGD) (Puillandre et al. 2012), Poisson Tree Processes (PTP) (Zhang et al. 2013) and General Mixed Yule-coalescent method (GMYC) (Pons et al. 2006). First, distance-based analysis ABGD was calculated using the K80 Kimura distance model and default parameters on the online platform (https://bioinfo.mnhn.fr/abi/public/abgd/abgdweb.html). Second, the single-locus species delimitation PTP method (Kapli et al. 2017) was used on the phylogeny excluding the outgroups *A. rufisignella* and *A. nummulalis*. Analyses were performed on the online platform (https://species.h-its.org). The confidence of delimitation schemes was assessed using MCMC chain of 10 million generations, a thinning of 100 and burn-in of 10% and the partition with the best likelihood were kept. Third, single threshold GMYC method was applied using the splits R package. The ultrametric tree required was generated using BEAST 1.10.4 (Suchard et al. 2018) with a GTR+G+I model, an uncorrelated relaxed clock with a lognormal distribution and a mtDNA COI substitution rate estimates of 0.0115 (Brower 1994). The species delimitation based on morphology has been compared to the delimitations based on molecular data. From the species described, the genetic p-distance was estimated between all pairs of samples using MEGA (Tamura et al. 2021) and summarized by species.

## Results

### DNA recovery from historical samples

Historical DNA extraction showed different yields mainly related to the age of the specimens but also to the type of tissue and the extraction method, i.e., destructive or non-destructive. Indeed, for the oldest sample, i.e., CRA03, captured between 1784-1789, it has been possible to extract DNA using a destructive approach on only one leg. For the two 19th century samples, i.e., CRA01 and CRA02, a non-destructive approach was attempted. These two samples are quite close in age, the large difference observed in extracted DNA between the two samples seems to be related to the tissue used and the extraction from the abdomen increases considerably the amount of extracted DNA. Surprisingly, the sample with the lowest concentration of DNA is the CRA07 sample which was captured in 1911 and for which two legs were used destructively. The size of the DNA fragments captured is also related to the age of the samples with smaller fragments for the sample captured during the 18th century, CRA03, and larger fragments for the sample CRA04 captured in 1992. Sample CRA07 is a special case since it was captured in 1911 but has small DNA fragments (Table 1). The specific degradation of this sample, low amount of DNA and small DNA fragments, may be related to different factors that we ignore including the conditions of capture and preservation.

For the more recent samples, captured after 2000, it was possible to extract enough DNA to perform a classical COI barcode amplification and Sanger sequencing (PCR amplification in Table 1). For the older samples, it has been necessary to develop a barcode capture approach because the amount of endogenous DNA was too low and initial trials at PCR amplification proved unsuccessful. This capture approach using probes designed along the COI barcode allowed NGS sequencing of 3.99 million reads in total with high heterogeneity between samples (Table 1). The number of reads seems correlated with the amount of DNA initially extracted. This heterogeneity in the amount of sequence recovered is then found throughout the bioinformatics analysis process until it impacts the percentage of barcode finally recovered. However, the capture approach was effective since it allowed the recovery of a sufficient proportion of barcode to perform phylogenetic inferences for each of the samples, including the oldest sample, CRA04, and the particularly degraded one CRA07 for which respectively more than 50% and 93.1% of the barcode was recovered.

### Phylogenetic inference and species delimitation

Phylogenetic inference has been performed on the whole COI barcode alignment including BOLD sequences, barcodes amplified for this study and sequences recovered using historical DNA capture approach. The samples corresponding to the two species *A. rufisignella* and *A. nummulalis* have been used as outgroups and their node is well supported (Figure 1). Then a well supported node separates two clusters, one includes the species *A. diplomochalis, A. insons* and *A. centrifugens* on one side and the species *A. gonogramma* and *A. lactella* on the other (Figure 1). Overall, the nodes separating the five species are also well supported. The three species delimitation analyses are consistent with each other and with morphology. The five species described and identified morphologically are almost all found by the species delimitation approaches, only the separation between *A. gonogramma* and *A. lacteella* has not been found in the ABGD approach based on the levels of divergence between sequences. However, the node separating the two species is well supported. Analysis of genetic divergence (p-distance) between each pair of individuals within and between species shows contrasting levels of divergence (Figure 2). Within species, genetic divergence is low between 0.43% for *A. centrifugens* and 2.47% for *A. diplomocalis* for which we have only three samples. The distribution of genetic divergence then shows a gap with much higher values between samples belonging to different species and a percentage of divergence ranging from 5.17% to 11.90%. These results confirm the species identified using morphology and species delimitation approaches. Within each species the species delimitation approaches also identified additional separations mainly related to geographic divergences. The details of the divergences within species will be discussed next in the “Molecular diagnosis” section of each species.

**Figure 1.**
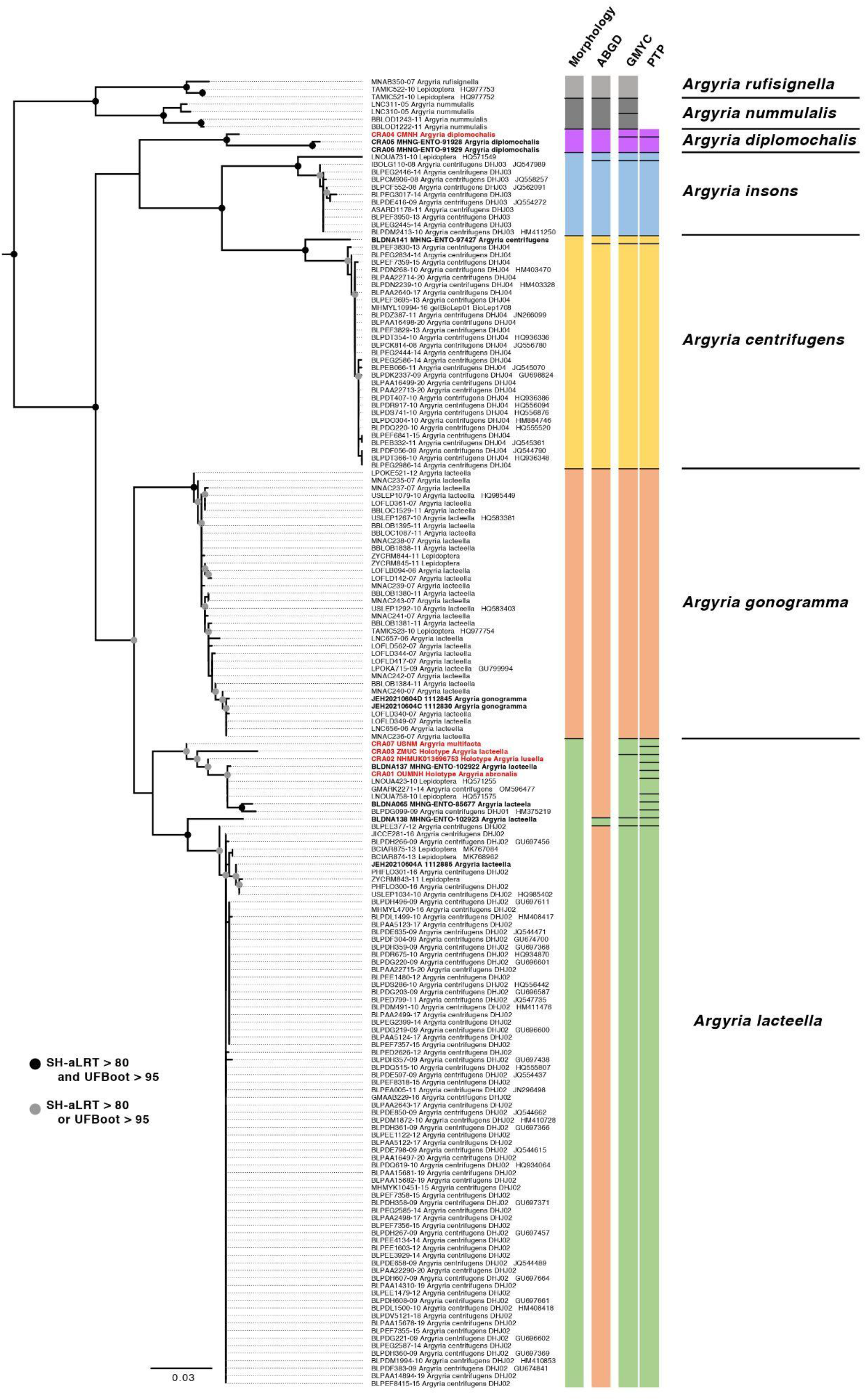
Phylogenetic inferences including 174 *Argyria* COI barcode sequences, i.e. 160 barcodes from BOLD, 9 COI sequences amplified by PCR (in bold), and 5 COI sequences obtained using capture from historical specimens (in red). Nodal support expressed in SH-aLRT and ultrafast bootstrap (UFBoot) is given as indicated in the caption except for nodes when SH-aLRT < 80 and/or UFBoot < 95. Species delimitation results including morphology, ABGD, GMYC and PTP are indicated by different colors to represent the species proposed.

**Figure 2.**
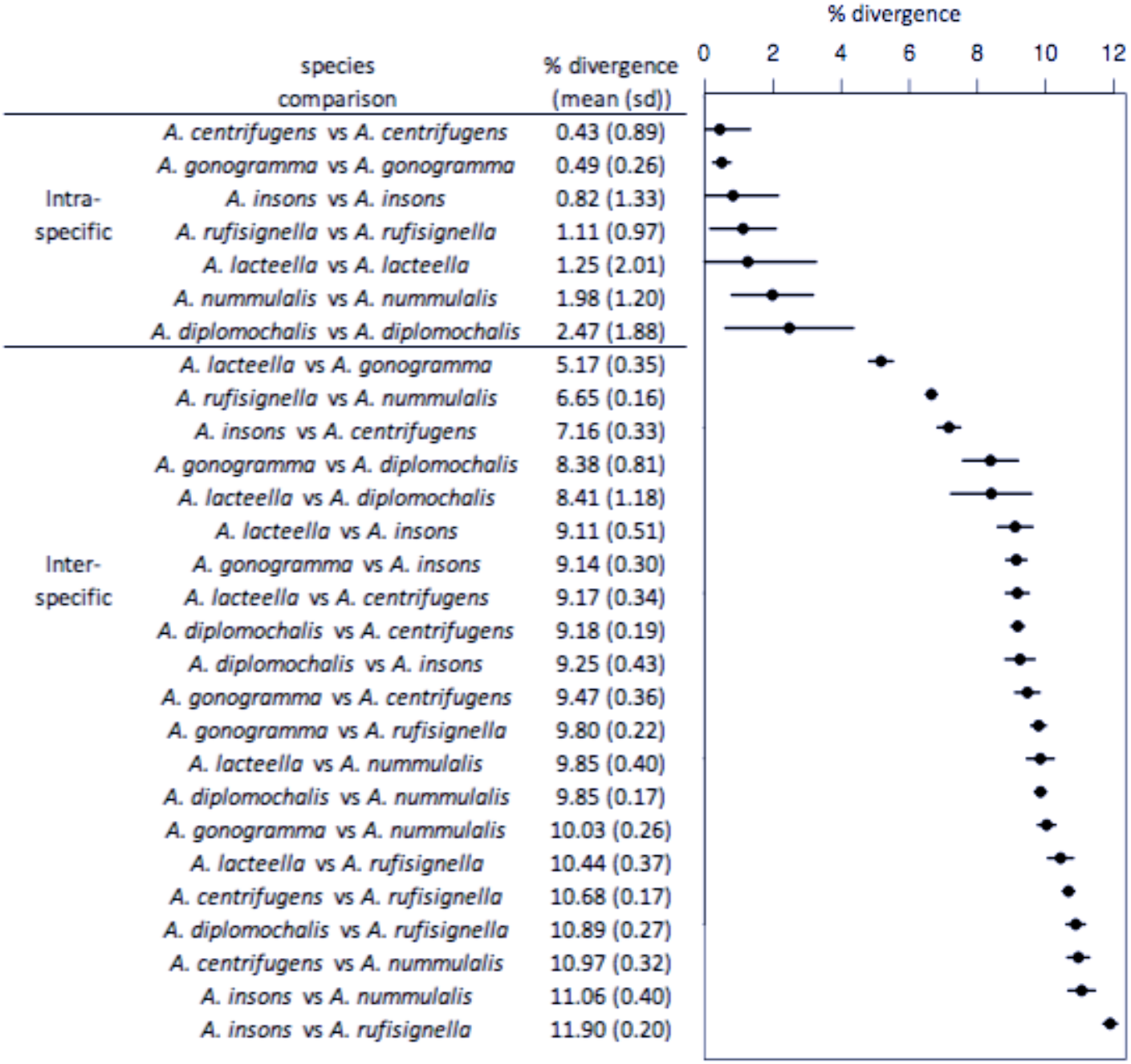
Mean genetic divergence (p-distance) between all samples from each species. The mean percentage of divergence is ordered according to intra or interspecific comparison.

### Taxonomic account

#### *Argyria lacteella* (Fabricius, 1794)

Figs 3-7, 11, 17, 24, 27, 29, 33

**Figure 3.**
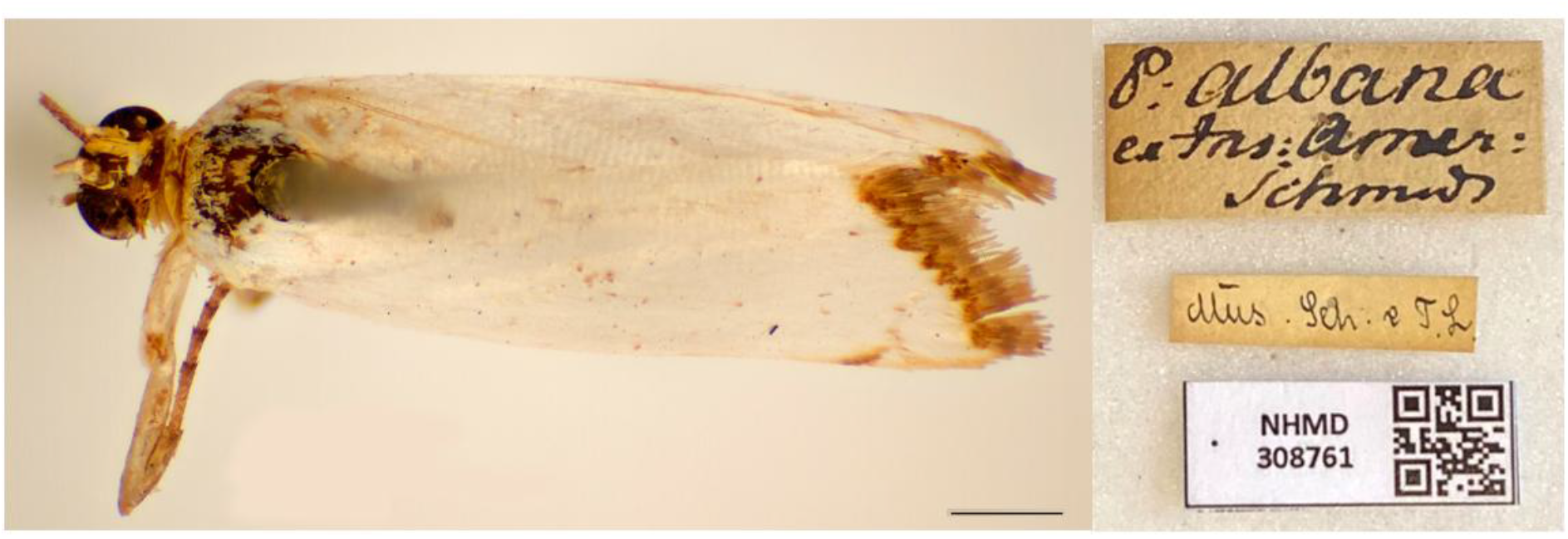
Holotype of *Tinea lacteella* Fabricius, 1794 (copyright of Natural History Museum of Denmark, ZMUC); scale bar: 1 mm.

*Tinea lacteella* Fabricius, 1794: 313. Type locality: “*Americae insulis*” (USA Virgin Island of Saint Croix; see Remarks).

- = *albana* (Fabricius, 1798 (*Pyralis*). Unnecessary replacement name.
- = *abronalis* Walker, 1859: 969 (*Zebronia*??). Type locality: Brazil, Rio de Janeiro.
- = *lusella* (Zeller, 1863: 51) (*Catharylla*). Type locality: St. Thomas Island [USA Virgin Islands]. **Rev. syn**.
- = *vestalis* Butler, 1878: 494, 495. Type locality: Jamaica. **Syn. nov**.
- = *multifacta* Dyar, 1914: 317. Type locality: Panama, Porto Bello. **Syn. nov**.

Fernald (1896: 72, plate V figs 4, 6); Dyar (1903: 411); Grossbeck (1917: 126; probably referable to *A. gonogramma*; see Remarks); Schaus (1940: 400); Amsel (1956: 31, pl. 69 fig. 6, part of records, misspelled ‘lactella’); Munroe (1956: 127); Błeszyński & Collins (1962: 214); Zimsen (1964: 579, misspelled ‘lactella’); Kimball (1965: 234, part of records); Błeszyński (1967: 96); Jaume (1967: 2); De la Torre y Callejas (1967: 20); Alayo & Valdés (1982: 61); Tan (1984: 96 *et seq*., misidentification); Ferguson et al. (1991: 40, misidentification); Munroe (1995a: 35); Heppner (2003: 288, part of the records); Martinez & Brown (2007: 81, fig. 9; referable to *A. gonogramma*); Roque-Albelo & Landry (2010); Scholtens & Solis (2015: 54); Landry et al. (2020: 101, fig. 6B); Gibson et al. (2021: 29).

#### Type material examined

**Lectotype** of *Tinea lacteella* (Fig. 3), **here designated**, with label data as follows: 1- “P. albana | ex Ins: Amer: | ?Schmud?”, 2- “Mus[eum]. S[ehested] & T[oender] L[und], 3- “LECTOTYPE | *Tinea lacteella* | Fabricius, 1794 | Des[ignated] by B. Landry, 2021”; deposited in ZMUC.

**Holotype** of *Zebronia? abronalis* (Figs 4, 24), with label data as follows: 1- “Type”, 2- “Rio”, 3- “91”, 4- “Zebronia | Abronalis”, 5- “TYPE LEP :No 1195 | Zebronia ? | abronalis | Walker | HOPE DEP[ARTMEN]T.OXFORD”; deposited in OUMNH.

**Holotype** of *Catharylla lusella* (Fig. 5), with label data as follows: 1- “Type”, 2- Lusella | Zell[er]. Mon[ograph]. p.51.”, 3- Zell[er]. Coll[ection]. | 1884.”, 4- “♂ | Pyralidae | Brit.Mus. | Slide No. | 7092” | DNA voucher Lepidoptera B. Landry, n° 00158 | NHMUK013696754 | MOLECULAR 215427977; deposited in the NHMUK.

**Holotype** of *Argyria vestalis* (Fig. 6), with label data as follows: 1- “Type”, 2- “Jamaica | 78. 19”, 3- ♂ | Pyralidae | Brit.Mus. | Slide No. | 7093 | NHMUK013696753”; deposited in the NHMUK.

**Holotype** of *Argyria pusillalis* variety *multifacta* (Fig. 7) with label data as follows: 1- “PortoBello | Pan[ama]. Febr[uary]. [19]11 | AugustBusck”, 2- “Type | No.16316 | U.S.N.M.”, 3- “Platytes | multifacta | Type Dyar”, 4- “♀ genitalia | slide 3826 | R W Hodges”, 5- “Genitalia Slide | By RWH ♀ | USNM 10,709”; deposited in the NMNH. **Paratypes** of *Argyria pusillalis* variety *multifacta* with label data as follows: 1 ♂: 1- “PortoBello | Pan[ama]. Febr[uary]. [19]11 | AugustBusck”, 2- “♂ genitalia | slide, 29 Apr. ‘32 | C.H. #29 | Genitalia slide | By ME ♂ | USNM 99,668”; 1 ♀: same data; 5 ♀♀, 2 ♂♂: same data except “Mar[ch]”; 2 ♀♀: 1- “RioTrinidad | Mar[ch]. [19]12 Pan[ama] | ABusck | coll”; 1 ♂: 1- “CorazolC[anal]Z[one] | Pan[ama] 3/24 [19]11 | AugBusck”, 2- “♂ genitalia | slide, 9 June. ‘32 | C.H. #83” [slide not found]; deposited in the NMNH. [Note: the Tabernilla (Busck) and Corazol (Crafts) specimens were not found at NMNH]

Other specimens examined: 238 specimens (see Supplemental material).

#### Morphological diagnosis

This is a small satiny white moth of 9.5-14 mm in wingspan. The forewing brown markings are median triangles on the costa and dorsal margin is usually linked by a thin straight line sometimes slightly thicker on the discal cell as a spot, but sometimes inconspicuous, another triangle subapically on costa, usually separated by a thin white line from a short oblique dash anteriorly, and a wavy terminal line (Figs 3-6, 11). There are also specimens of *A. lacteella* with a complete median fascia (Fig. 7) in the South of the distribution of the species, in Panama (holotype of synonym *A. multifacta*), French Guiana, Bolivia, and Brazil. In forewing markings *A. lacteella* differs from *A. gonogramma* (Figs 8, 12), which usually has a well-marked darker, blackish-brown spot on the discal cell, linked by a thin, curved line to a short diagonal bar on costa and a thin triangle on the dorsal margin, and without a clear costal triangle subapically. In forewing markings *A. lacteella* is most similar to *A. centrifugens* (Fig. 10), which is generally bigger (14-19 mm in wingspan) and which has the line anteriad to the subapical costal triangle curved to reach the costa at right angle whereas that line in *A. lacteella* runs obliquely into the costa. In male genitalia (Fig. 17) this species differs from the most similar *A. gonogramma* by the basal projection of the valva that is slightly longer and bent mesad at right angle whereas it is just barely curved in *A. gonogramma* (Figs 16, 18). The cornuti on the vesica also are smaller and thinner in *A. lacteella* compared to those of *A. gonogramma*. In female genitalia *A. lacteella* (Figs 24, 27) is also most similar to *A. gonogramma* (Fig. 28), but *A. lacteella* has two “pockets” anterolateral of the ostium bursae, whereas *A. gonogramma* has one continuous pocket anterior of the ostium.

**Figures 4-7.**
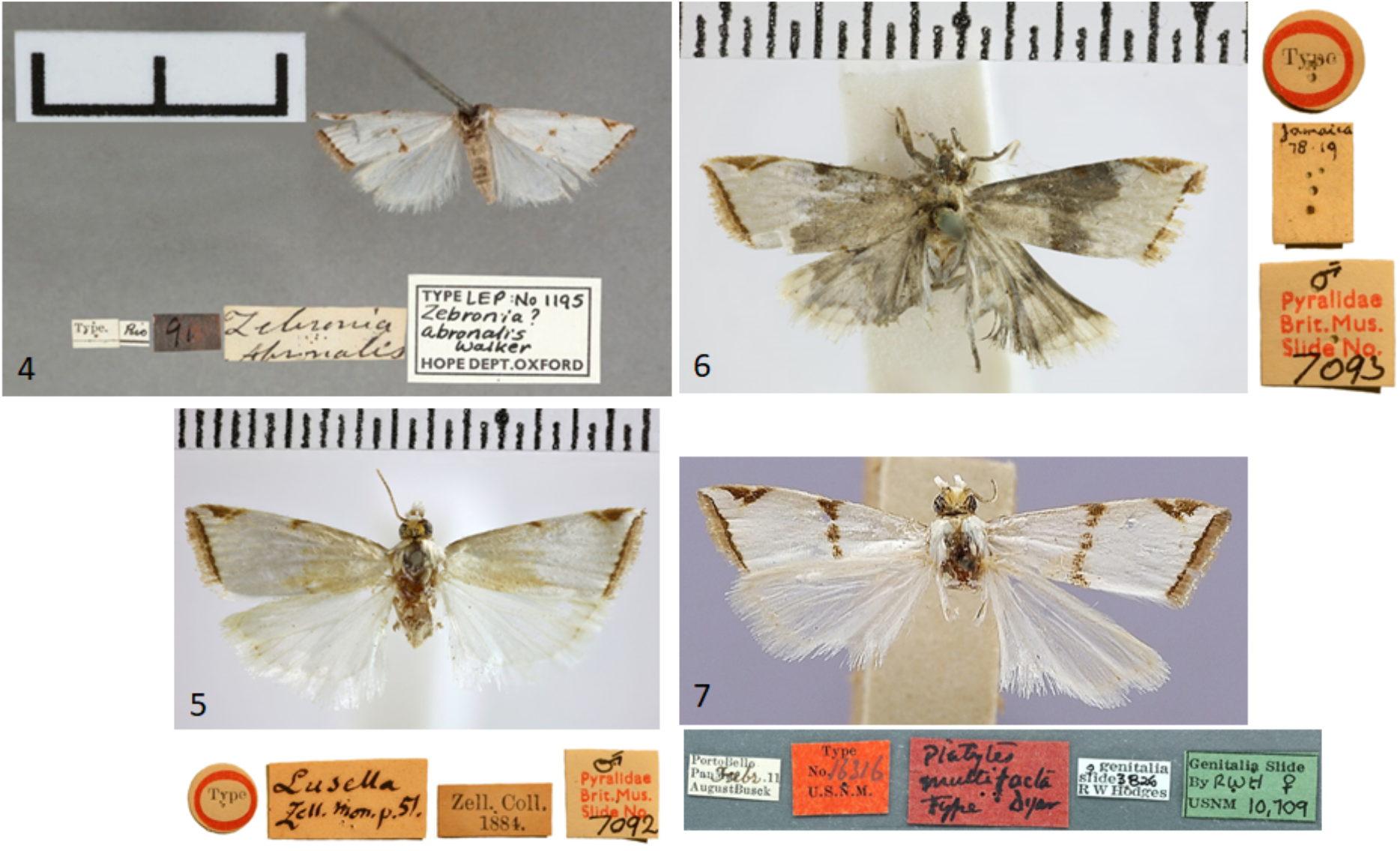
Holotypes of *Argyria lacteella* synonyms **4** *Zebronia*?? *abronalis* Walker, 1859 (© Oxford University Museum of Natural History; OUMNH); scale bar: 10 mm) **5** *Catharylla lusella* Zeller, 1863 (NHMUK © Trustees of the Natural History Museum) **6** *Argyria vestalis* Butler, 1878 (NHMUK © Trustees of the Natural History Museum) **7** *Argyria pusillalis* variety *multifacta* Dyar, 1914 (NMNH; wingspan: 12 mm).

#### Molecular results

Phylogenetic inference based on COI barcode alignment reveals a large cluster grouping the *A. lacteella* samples. This clade is relatively homogeneous since the percentage of divergence within this species remains low with an average of 1.25% (Fig. 2). It is then divided into 2 clusters also identified by the species delimitation approaches. The first one is mainly composed of samples from South America, i.e. Brazil, French Guiana, Argentina, Colombia, but also from the Galapagos Islands. The historical samples originating from Panama and the United States Virgin Islands belong to this clade. The different intraspecific delimitations identified by the species delimitation approaches within this clade are therefore certainly related to geographical divergence. The second cluster is composed of a clade of samples from the US on one side and a very large clade of samples from Costa Rica on the other. The latter shows a very low level of variation.

#### Distribution

Fig. 33. Widespread in the Western Hemisphere from the USA State of Florida north to Alachua County in the north, across Central America and the Antilles, in South America to Argentina in the south, as well as on the Galápagos Islands.

#### Remarks

Fabricius (1798) changed the name of his *lacteella* (1794) with another (*albana*). The reason for this is unrecorded and remains unclear, but this is possibly because Fabricius (1798: 476, spelling it “*lactella*”) incorrectly considered *lacteella* to be a homonym of *Tinea lactella* Denis & Schiffermüller, 1775 (a synonym of *Endrosis sarcitrella* (Linnaeus)). Also, he may have corrected ‘improper’ names as in his treatments of *Tinea compositella* Fabricius, 1794 and *T. tapetzella* Linnaeus, 1758, both without putative ‘homonyms’ and respectively renamed *Pyralis composana* and *P. tapezana* (Fabricius, 1798: 480), or he felt thought that the exact orthography of any name was not so important.

The first label associated with the lectotype of *A. lacteella* (Fig. 3) reads “P[yralis]. albana | ex Ins. Amer: | Schmidt”. The second line of this label means “from the American Islands” while the name of the third line refers to the collector of the specimen, who, according to Zimsen (1964) was either Adam Levin Smidt, a custom-house officer, or Johan Christian Schmidt, a surgeon. Both lived on the island of St Croix, which was at that time a Danish possession (T. Pape, pers. comm. to BL). The second label associated with this type specimen refers to the collection of Ove Ramel Sehested and Niels Tønder Lund who lived in Copenhagen and were pupils and friends of Fabricius (Baixeras & Karsholt, 2011).

The locality of origin is an additional complication associated with *A. lacteella*. The locality of Fabricius’ *Pyralis albana* (1798) is mentioned as “*Americae insulis*” [American islands] whereas that of *A. lacteella* (1794) is “*Americae meridionalis arboretis*” [South American arboretum]. Given that “Dr. Pflug” is mentioned in the original description of *A. lacteella*, it is reasonable to conclude that “*Americae insulis*” was a correction for “*Americae meridionalis arboretis*”. This is because Paul Gottfrid Pflug (1741-1789), a medical doctor, lived in the Caribbean island of Saint Croix (United States Virgin Islands) during the last five years of his life, where he collected insects that he sent to Denmark. He is mentioned often by Fabricius as a specimen collector (O. Karsholt, pers. comm. to BL, 3.vi.2021). Therefore, *A. lacteella*/*albana* is from an American island (*Americae insulis*) that is probably Saint Croix. As confirmed by Copenhagen Museum former curator Ole Karsholt and present curator Thomas Pape, only one type specimen presently exists for *lacteella*/*albana* (Fig. 1) and because *albana* is best considered as an unjustified replacement name, the type of *Pyralis albana* Fabricius, 1798 is the same as that of *Tinea lacteella* Fabricius, 1794. This specimen is without an abdomen and is designated as the lectotype upon the recommendation of curator T. Pape, who wrote (7.vi.2021 to BL): “As Fabricius does not indicate the number of specimens, I would consider a lectotype designation as appropriate, unless this has already been done by referring to this specimen as “the type” or something similar.” Such a designation also serves to stabilize the identity of the species name laden with confusion caused by Fabricius himself.

The type specimen of *A. lacteella* (Fig. 3) is badly rubbed, lacking most scales on the head, and some on the thorax and forewings as shown by the denuded anal vein on the right forewing. It also lacks most of the diagnostic brown markings of the forewing, notably the subapical triangle on the costa, but the terminal zigzagging brown line is almost complete and there are a few brown scales in the position of the median spot and fewer brown scales still on the dorsal margin medially.

Munroe (1995: 35) stated that *Argyria abronalis* is a *nomen dubium*, but this is incorrect as a female type (Figs 4, 24) is in the Oxford University Museum of Natural History (also figured at https://www.oumnh.ox.ac.uk/collections-online#/item/oum-catalogue-3393). The forewing markings, female genitalia morphology and DNA barcode all concur to validate the synonymy of *A. abronalis* with *A. lacteella*. The species was described from the female sex, without indication or indirect evidence of more than one specimen; therefore the OUMNH specimen is considered the holotype.

The original description of *Catharylla lusella* Zeller (1863) explicitly mentioned one female only, described from the island of Saint Thomas, US Virgin Islands. Thus, the male sign on this holotype’s slide number label is incorrect (Fig. 5). The specimen was dissected and although the genitalia dissection was not thoroughly cleaned, the visible morphological characters agree with those of *A. lacteella*. The forewing markings lack the median triangle of the dorsal margin and any indication of a median transverse line, as in the holotype of *A. lacteella*, but the subapical triangle on the costa and especially the COI barcode obtained clearly show that *C. lusella*, **rev. syn**., should be considered a synonym of *A. lacteella*. The name had been synonymized by Fernald (1896), considered a synonym also by Schaus (1940), but considered valid again by Błeszynski & Collins (1962) and Munroe (1995; misspelled “*lusalla*”).

The original description of *Argyria vestalis* does not mention more than one specimen and the NHMUK does not hold additional specimens with these label data; therefore, this specimen is considered the unique holotype. It is a lightly marked, damaged, dissected male (Fig. 6); the dissection clearly shows the curved projection at the base of the valva that is diagnostic for *A. lacteella*. Therefore the name *A. vestalis* **syn. nov**. is considered a synonym of *A. lacteella*.

*Argyria multifacta* was described as a variety of *A. pusillalis* for which “All the specimens have the median band continuous across the wing” (Dyar, 1914: 317). Among a series of specimens mentioned from several localities in the Panama Canal zone, one is recorded as Type with the type number and label data mentioned above (Fig. 7). Although this holotype shows a conspicuous and almost continuous median band on the forewing, thus revealing strong variation in that respect in the species, other wing characters, size, and the COI barcode data point to the synonymy of *A. multifacta* **syn. nov**. with *A. lacteella*.

This species evidently became established in Florida, USA in the 1970s and consequently, earlier records from Florida (Grossbeck, 1917; Kimball, 1965) are believed to be wrong and referable to *A. gonogramma*. The earliest specimen known to us was collected in Miami-Dade County, Fuchs Hammock near Homestead, by T.S. Dickel on 31 August 1979 (MGCL catalog no. 1112898, slide 6219, deposited in FSCA). The species rapidly spread across the state, as shown by first collection years in other vouchered counties: 1983: Highlands, Monroe, Orange; 1986: Collier, Manatee; 1987: Volusia; 1988: Lee; 1990: Pinellas; 1991: Hernando; 2000:

Brevard; 2003: Marion; 2005: Alachua; 2012: Indian River; 2013: Levy (FSCA, MGCL). The collection of *A. gonogramma* in Florida decades before *A. lacteella* strongly suggests that the latter species is non-native and that it invaded in the given time frame (see Remarks for *A. gonogramma*).

Amsel (1956: 31, pl. 69 fig. 6,) mentions the species from specimens sporting a wingspan of 12 to 18 mm, and although his illustration probably represents *A. lacteella*, no specimens examined of that species were found to reach a wingspan of more than 14 mm.

#### *Argyria gonogramma* Dyar, 1915

Figs 8, 12, 13, 15, 16, 18, 28, 30, 34

*Argyria gonogramma* Dyar, 1915: 87–88. Type locality: Bermuda.

= *pusillalis* Hübner, 1818: 28, Figs 167, 168. Type locality: [USA, Maryland] Baltimore. *Nomen dubium*.

= *pussillalis* Hübner, 1818: 28. Original misspelling.

Bleszynski & Collins (1962: 213).

Recorded as *Argyria lacteella* (Fabricius, 1794): Fernald (1896: 72, plate V fig. 5); Grossbeck (1917: 126); Kimball (1965: 233); Tan (1984: 96 *et seq*.); Ferguson et al. (1991: 40); Munroe

(in part, 1995: 35); Martinez & Brown (2007: 81, fig. 9).

#### Type material examined

**Holotype** ♂ (Figs 8, 15, 16), with label data as follows: 1- “Bermuda, | 11.3.BWI | F.M. Jones”, 2- “V-3 | D”, 3- “Type No. | 18244 | U.S.N.M.”, 4- “Argyria | gonogramma | Type Dyar”, 5- “♂ genitalia | slide, 29Apr[il].’32 | C.H. #27”, 6- “Genitalia Slide | By 107,454 | USNM”; deposited in the NMNH.

Other specimens examined: 411 specimens (see Supplemental material).

#### Morphological diagnosis

In this small satiny-white moth measuring between 10.5 and 13.5 mm in wingspan, the median markings of the forewing (Figs 8, 12) usually include a well-marked blackish-brown spot on the discal cell that is connected by curved lines to an oblique bar on the costa and a thin triangle on the dorsal margin. On the forewing costa, subapically, a thin curving bar is not followed by a triangle, but usually by 1-3 horizontal lines reaching the terminal margin below the apex. Relatively dark brown forms, with less contrasting markings (Fig. 13) have been collected in Alabama, Florida, and Louisiana (CUIC, FSCA, MHNG, NMNH) from December to April. In forewing markings this species is closest to *A. lacteella* (Fig. 11) in which there usually is a clear subapical triangle on the costa and for which the median spot, if present, is paler brown and usually smaller than the costal and dorsal triangles. In the absence of a subapical triangle on the forewing costa *A. gonogramma* is also similar to *A. diplomochalis* (Figs 9, 14), which, however, doesn’t have any indication of a median spot or of any line between the median spot of the dorsal margin and the costa. In the male genitalia *Argyria gonogramma* (Figs 15, 16, 18) has the basal projection of the valva shorter than that of *A. lacteella* (Fig. 17) and just barely curving (Fig. 16); the cornuti on the vesica are also longer and thicker than those of *A. lacteella*. In the female genitalia (Fig. 28) only one wide, sclerotized pocket can be found anterior of the ostium bursae, whereas *A. lacteella* (Figs 24, 27) has two pockets in the same area.

**Figures 8-10.**
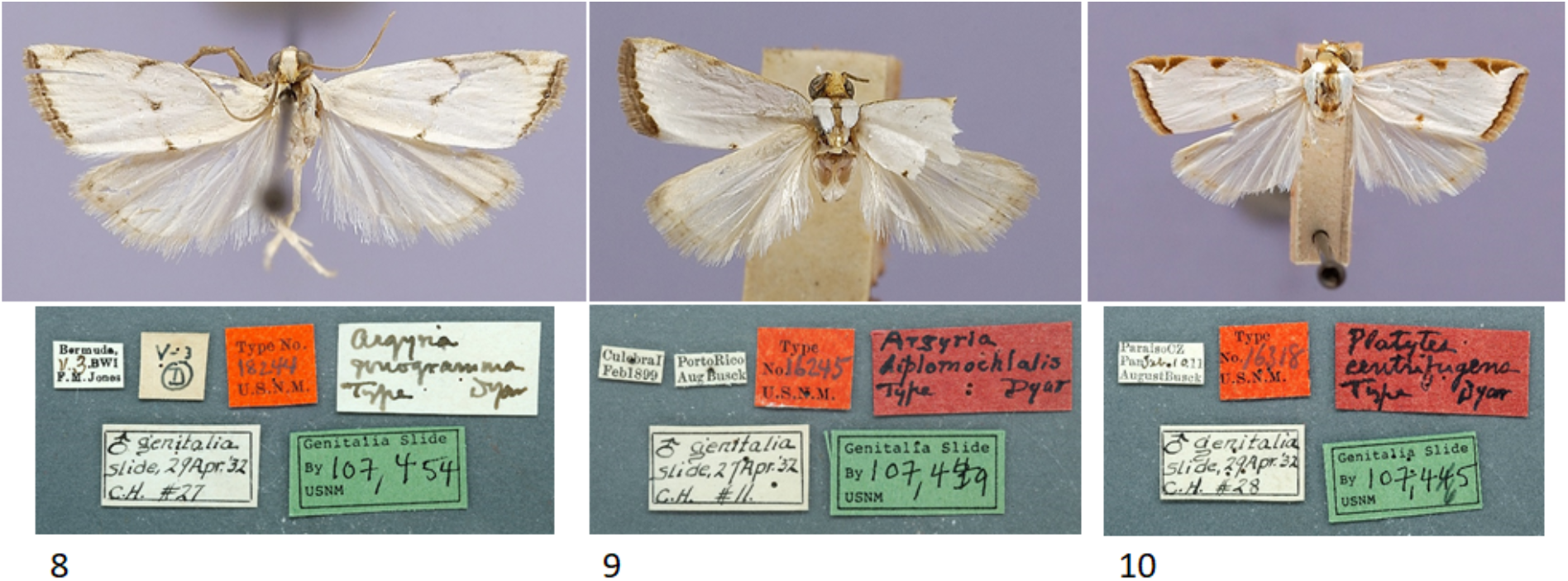
Habitus of *Argyria* type specimens (all in NMNH) with labels underneath **8** Holotype of *Argyria gonogramma* Dyar, 1915 (wingspan: 11 mm) **9** Lectotype of *Argyria diplomochalis* Dyar, 1913 (wingspan: 11 mm) **10** Holotype of *Argyria centrifugens* Dyar, 1914 (wingspan: 16 mm).

#### Molecular results

Phylogenetic inference reveals that *Argyria gonogramma* constitutes a homogeneous clade. The monophyletic clade is identified in both GMYC and PTP species delimitation approaches, but it is not found in the ABGD approach and is grouped with *A. lactella* (Fig. 1). This clade shows very low genetic variability within the COI barcode with an average intraspecific divergence of only 0.49% (Fig. 2). This low genetic diversity may be the result of different evolutionary processes, including recent colonization. This species is mainly present in the US where it overlaps with *A. lacteella* in Florida (Figs 33-34).

#### Distribution

Fig. 34. Bermuda, Bahamas, widespread in the Eastern USA, from North Carolina in the North to the south of Florida, west to eastern Texas.

#### Remarks

The specimen of *A. gonogramma* labelled ‘Type’ in the NMNH is considered the unique holotype; the species’ description (Dyar, 1915) doesn’t indicate multiple specimens. *Argyria pusillalis* Hübner is associated here with *A. gonogramma* and not with *A. lacteella* as in Munroe (1995) because at the latitude of Baltimore, Maryland, U.S.A., the type locality of *A. pusillalis*, only the superficially similar *A. gonogramma* or *A. rufisignella* (Zeller, 1872) could occur. *Argyria nummulalis* Hübner, 1818 is also known to occur in the eastern USA at the latitude of Baltimore, but this species lacks any median markings across the forewing, unlike the illustration of *A. pusillalis*. The name *A. pusillalis* is considered a *nomen dubium* because the original description and illustration associated with it do not allow a conclusive determination. Hübner’s collection was deposited in the Naturhistorisches Museum Wien, Vienna, Austria, which was destroyed by fire in 1835 (Horn et al., 1990). However, although type specimens of some of Hübner’s Noctuidae species have recently been discovered in this museum (Gabor and Laszlo Ronkay, pers. comm. to BL), a search for a type specimen of *A. pusillalis* was not successful (S. Gaal, pers. comm. to BL). This issue could be settled by the designation of a neotype, but we refrain from doing that in order to avoid more instability in the nomenclature of this group. Also, we believe that it should be done in conjunction with a taxonomic revision of *A. rufisignella*, at the least.

*Argyria pusillalis* was originally named “*pussillalis*” (Hübner, 1818: 28), then mentioned as “*pusillalis*” on page 30 and on two indices (pages [36] and [38]), and finally as “*pussillalis”* again on the plate with the illustrations. Given that “*pusillus*” is Latin for small, it seems reasonable to believe that the original spelling “*pussillalis*” was in error.

*Argyria gonogramma* is a North American native species that was previously misidentified as *A. lacteella* and that has been collected in the United States since the late 1800’s. The earliest specimens in the NMNH were collected by C.V. Riley from Ar[t]elier, FL, 1882, and an undated specimen from N.[orth]C.[arolina]. Another specimen collected by Boll in Texas (collection date unknown) was identified by “Rag[onot] \[18]86”, and then by “CVR[iley]at the B. Mus. \[18]87”.

That this species is native to the Southeastern U.S., or at least was established long before *A. lacteella*, as shown by earlier collecting dates for specimens in the FSCA and MGCL. For example: Florida, Sarasota Co.: 1951, Alachua Co.: 1960, Volusia Co.: 1962, Okaloosa Co.: 1963, Texas: 1978, Louisiana: 1979.

The earliest record of *A. lacteella* in 1979 in the USA (Florida) supports the conclusion that Tan (1984), although referring to *A. lacteella*, was in fact dealing with *A. gonogramma*. Tan’s (1984) unpublished MSc thesis provided a description of larvae, with setal maps, which were reared from egg to adult on St. Augustine grass (*Stenotaphrum secundatum* (Walt.) Kuntze; Poaceae). Tan (1984) further mentions that the early instar larvae eat the upper epidermis only and when not feeding, larvae hide in shelters made of leaves attached with silk, wherein molting occurs. Tan (1984) also records the construction by the mature larva of a “small, compact silken case covered with frass and tiny pieces of chewed grass for pupation.”

Based on collected series of specimens both *Argyria gonogramma* and *A. lacteella* now occur in sympatry and fly on the same dates in Florida, for example at Archbold Biological Station in Highlands County or in Pinellas County.

Fernald (1896) treated this species under *A. lacteella* (pl. V fig. 5), whereas his other illustrations on the same plate (Figs 4 and 6) represent the true *A. lacteella*. Grossbeck (1917), Kimball (most or all records, 1965), and Martinez & Brown (2007) all treated this species under *A. lacteella*. Melanic specimens collected in winter months account for the specimens of “*A. diplomochalis*” cited by Kimball (1965). This colouration variant may represent an adaptation for hiding in dry grass during the winter months and/or to obtain extra calories from the sun to allow biological activity.

The single moth at the basis of the Vermont record has been dissected and is correctly determined, but it is far outside the range since we know of no other record of *A. gonogramma* north of North Carolina. It was collected in sandplain habitat (M. Sabourin, pers. comm. to JH), which is consistent with the species’ habitat preference in Florida.

**Figures 11-14.**
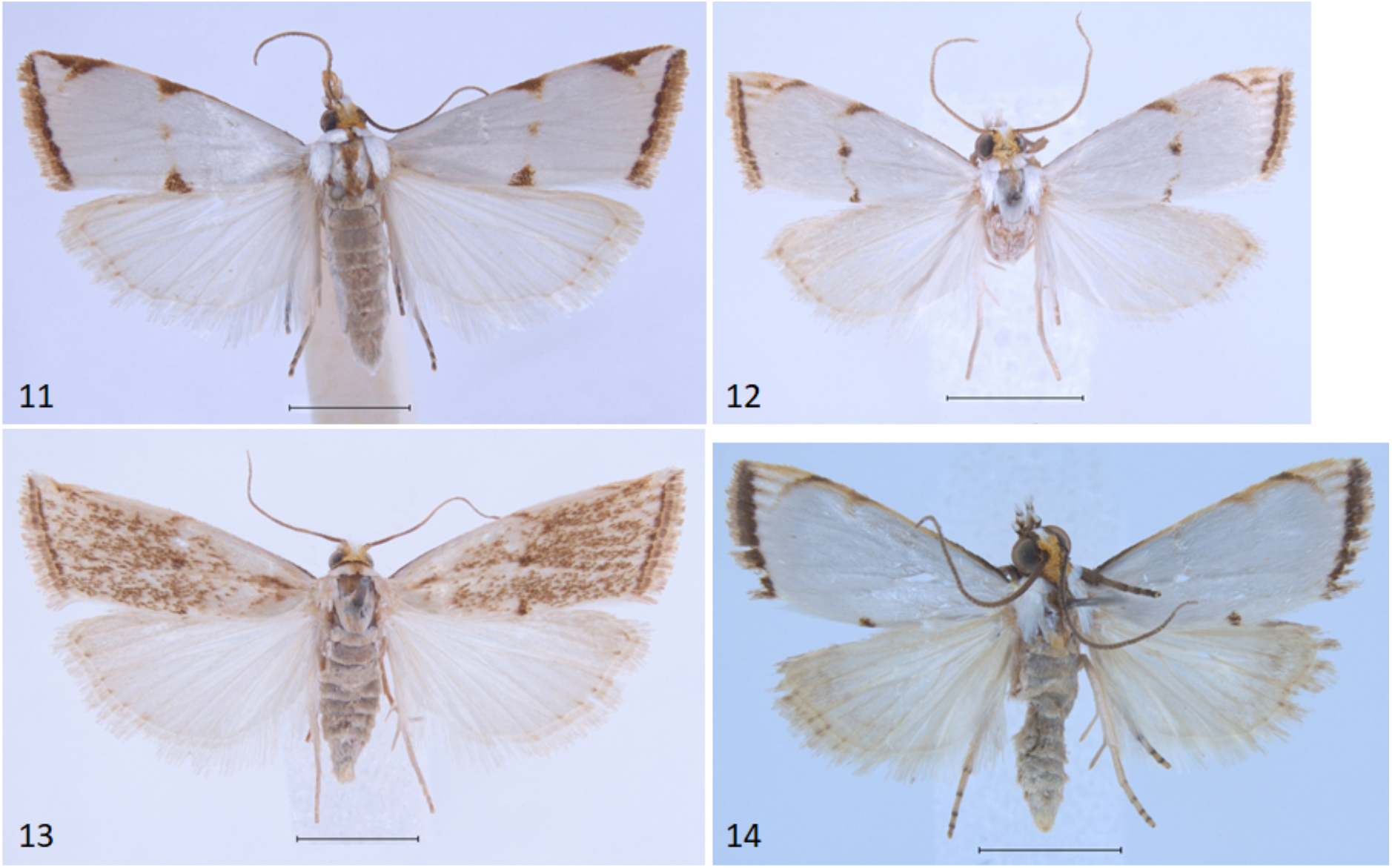
Habitus of additional *Argyria* specimens **11** *Argyria lacteella* (Florida, Putnam Co., FSCA) **12** *Argyria gonogramma* (Florida, Seminole Co., MHNG) **13** *Argyria gonogramma* (dark form, Louisiana, Calcasieu Co., MHNG) **14** *Argyria diplomochalis* (St Croix Island, MHNG). Scale bars: 2.5 mm.

**Figures 15-16.**
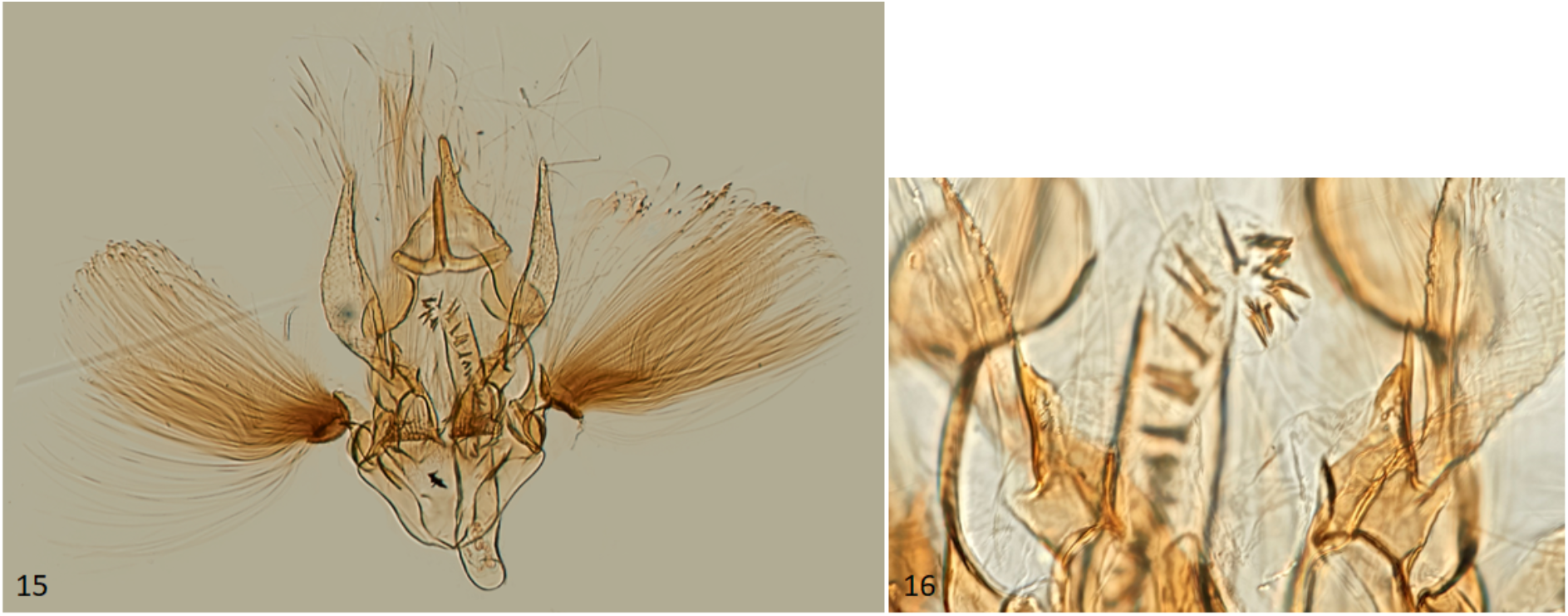
Male genitalia of *Argyria gonogramma* holotype (NMNH) **15** Whole genitalia **16** Close-up of bases of valvae with tip of phallus in middle.

**Figures 17-18.**
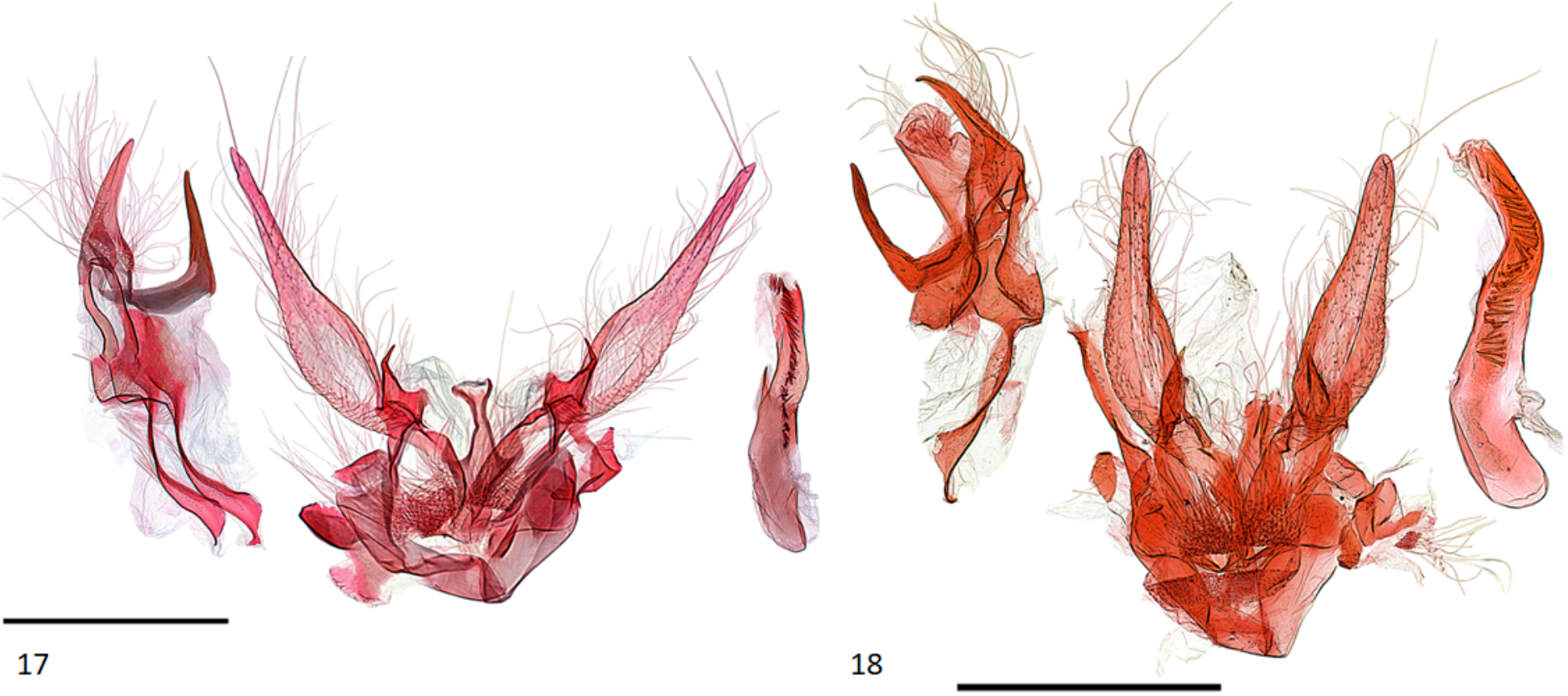
Male genitalia of *Argyria* species **17** *Argyria lacteella* (Florida, Alachua Co.) **18** *A. gonogramma* (Florida, Wakulla Co.). Both in FSCA; scale bars: 500 µm.

#### *Argyria diplomochalis* Dyar, 1913

Figs 9, 14, 19, 20, 25, 31, 35

*Argyria diplomochalis* Dyar, 1913: 113. Type locality: [USA] Culebra Island, Puerto Rico. Schaus (1940: 400, misspelled ‘diplamachalis’); Błeszyński & Collins (1962: 213); Kimball (1965: 234, US specimens misidentified); Błeszyński (1967: 96); De la Torre y Callejas (1967: 20); Alayo & Valdés (1982: 61); Munroe (1995: 35).

#### Type material examined

**Lectotype** ♂ (Fig. 9), here designated, with label data as follows: 1- “CulebraI[sland] | Feb[ruary]1899”, 2- “PortoRico | Aug[ust.] Busck”, 3- “Type | No.16245” | U.S.N.M.”, 4- “Argyria | diplomochalis | Type Dyar”, 5- “♂ genitalia | slide, 27 Apr[il].[19]’32 | C.H. #11.”, 6- “Genitalia Slide | By CH | 107,449 | USNM”, 7- “Lectotype | Argyria | diplomochalis Dyar, 1913 | Des[ignated] by M.A. Solis, 2022”, deposited in the NMNH. **Paralectotypes** (8 ♂, 1 ♀), here designated with label data as follows: 3 ♂♂: 1- “Culebra I[sland] | Feb[bruary]1899”, 2- “Porto Rico | Aug[ust] Busck”; 3 ♂♂: 1-”Bayamon | Jan[uary]1899”; 2- “Porto Rico | Aug[ust] Busck”; 1 ♀: 1-”Bayamon | Jan[uary]1899”, 2- “Porto Rico | Aug[ust] Busck”, 3-”GS-5620-SB | Argyria ♀ | lusella Z[eller] | det. Bleszynski, 19”, 4-”Genitalia slide | By SB ♀ | USNM 52861”; deposited in NMNH. 1 ♂ [abdomen in vial]: 1- “SYN-TYPE”, 2- “Bayamon | Jan 1899”, 3- “Porto Rico | Aug[ust] Busck”, 4- “Bleszynski | Collection | B.M. 1974-309”, 5- “Argyria | lusella Z[eller]. | ♂ | det. Bleszynski”, 6- “SYNTYPE | Argyria | diplomochalis | Dyar | det. M. Shaffer, 1975”, 7- “NHMUK013697137”; 1 ♂: 1- “SYN-TYPE”, 2- “Culebra I[sland]. | Feb 1899”, 3- “Porto Rico | Aug[ust] Busck”, 4- “Bleszynski | Collection | B.M. 1974-309”, 5- “SYN-TYPE” | Argyria | diplomochalis | Dyar | det. M. Shaffer, 1975”, 6- “NHMUK013697138”; deposited in the NHMUK.

Other specimens examined: 41 (see Supplemental material).

#### Morphological diagnosis

Measuring between 10 and 13 mm in wingspan this species (Figs 9, 14) is quite similar in size and forewing markings to *A. gonogramma* (Figs 8, 12), but it lacks a median line on the forewing and the median marginal markings are reduced to a faint brown bar on costa and a small dark-brown spot on the dorsal margin; the forewing costa and the head also appear more strongly marked (Fig. 31), notably more thickly dark brown at the base of the costa, on the frons laterally and on the labial and maxillary palpi; the costa of the forewing is also gold yellow to the apex, following the dark brown base. In male genitalia (Figs 19, 20), this species differs most noticeably from the others treated here in the thicker and less strongly bent apical section of the gnathos and in the short valva with a prominent sickle-shaped projection at its base. In female genitalia (Fig. 25), *A. diplomochalis* differs from the others treated here more noticeably by the long and narrow ductus bursae without sclerotized section as well as in the large, circular corpus bursae; the ostium region also lacks any sclerotized ‘pockets’.

**Figures 19-20.**
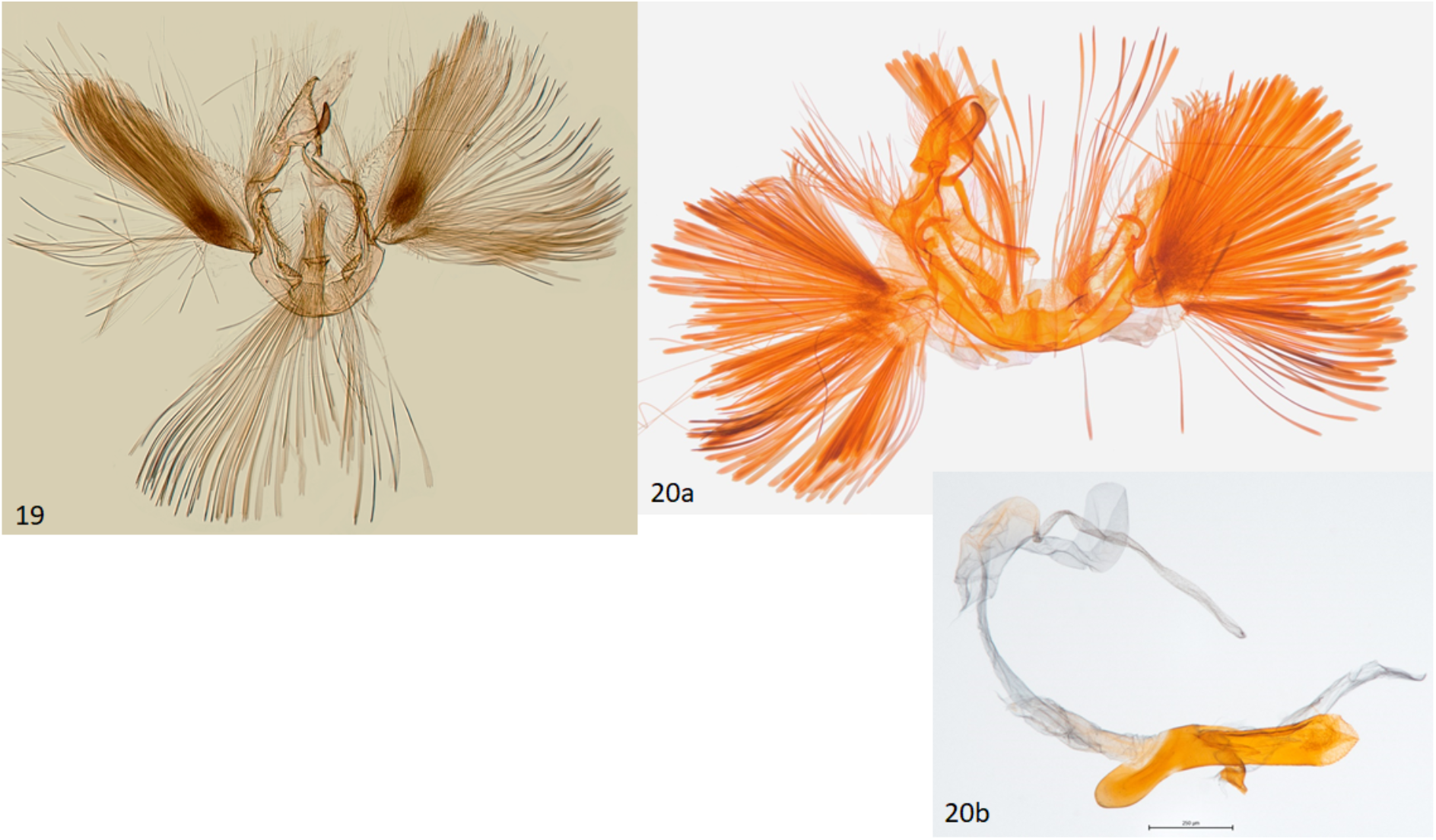
Male genitalia of *Argyria diplomochalis* **19** Holotype (NMNH) **20 a-b** Specimen from St Croix Island (MHNG-ENTO-91929) with phallus detached (**b**).

#### Molecular results

The phylogenetic clade corresponding to the species *A. diplomochalis*, comprises only three samples sequenced for this study. It appears that no sequence available in the BOLD database corresponds to this species. All three species delimitation approaches identify this clade (Fig. 1) and the average genetic divergence with the two closest species, *A. insons* and *A. centrifugens*, are respectively 9.25% and 7.16%, which confirms the high divergence of this clade and its specific status. Within this species, the GMYC and PTP approaches separate CRA04 on one hand and CRA05 and CRA06 on the other; and a mean divergence of 2.87% is observed between the samples of this clade. But this divergence may be related to a geographical divergence since CRA04 comes from the island of Anguilla and CRA05-CRA06 come from the US Virgin Island of Saint Croix. The integration of a larger number of samples in a genetic study could allow a finer molecular characterization of this species.

#### Distribution

Fig. 35. Antilles, from Cuba in the West to Dominica in the Lesser Antilles in the east.

#### Remarks

Described from 12 cotypes from Culebra Island and Bayamon, Puerto Rico, a lectotype is designated here to ensure that the name continues to refer to this species exclusively. Dyar (1913) stated “Cotypes, 12 specimens”, but only seven specimens were found at the NMNH while two others (now paralectotypes) are in the NHMUK.

Examination of specimens of “*A. diplomochalis*” cited by Kimball (1965), including ones in the FSCA labeled “5958,1” (Kimball’s number for that species), are *A. gonogramma* with scattered honey-brown scales on the forewings. This may be melanism caused by pupation during cold weather; all the specimens have been collected in winter or early spring.

#### *Argyria centrifugens* Dyar, 1914

Figs 10, 21-23, 26, 32, 35

*Argyria centrifugens* Dyar, 1914: 318. Type locality: Panama, Canal Zone, Paraiso. Błeszyński & Collins (1962: 212); Błeszyński (1967: 96); Munroe (1995: 35); Miller et al. (2012: 11); Landry et al. (2020: 101, fig. 6A);

#### Type material examined

**Holotype** ♂ (Figs 10, 21, 22), with label data as follows: 1- “ParaisoC[anal]Z[one] | Pan[ama]. Febr[uary].10.[19]11 | AugustBusck”, 2- “Type | No.16318 | U.S.N.M.”, 3- “Platytes | centrifugens| Type Dyar”, 4- “♂ genitalia | slide, 29Apr[il].[19]’32| C[arl].H[einrich]. #28”, 5- “Genitalia Slide | By 107,465 | USNM”; deposited in the NMNH.

Other specimens examined: 87 specimens (see Supplemental material).

#### Morphological diagnosis

*Argyria centrifugens* (Fig. 10) is very similar in wing markings to *A. lacteella* (Fig. 11), with which it can occur in sympatry in Central and South America, although the median line is always thin and not more pronounced in the middle or wide as in some South American specimens of *A. lacteella* (Fig. 7). It is also a bigger species, sporting a wingspan of 16 (male holotype)-17 mm in males and 16-19 mm in females, compared to 9.5- 12.0 mm in males and 11.0-14.0 mm in females of *A. lacteella*. Apart from size these two species differ in the colouration of their labial palpi as those of *A. lacteella* (Fig. 29) and *A. gonogramma* (Fig. 30) are pale greyish brown and yellowish gold with the apex satiny white whereas those of *Argyria centrifugens* (Fig. 32) are mostly dark brown with paler scales on the first palpomere but with the third palpomere dark brown to slightly paler brown. Both species are also very different in genitalia. The male genitalia of *A. centrifugens* (Figs 21-23) differ most notably in the three-pronged gnathos, the wider valva with a widely rounded apex and without a short hook-like projection at base but with a large membranous structure sporting a thin and pointed rod about half as long as the valva, directed toward the base of the valva and apparently articulated. The entire female genitalia are about twice as long in *A. centrifugens* (Fig. 26) than in *A. lacteella* (Fig. 24), the ostium is surrounded by a broad chamber with sclerotized wrinkles on the ventral wall, and the ductus bursae at the base is a medium-sized, thickly sclerotized tube in *A. centrifugens* whereas in *A. lacteella* the antrum consists of two lateral pockets of medium size and the base of the ductus bursae is a lightly sclerotized and striated round pocket.

#### Molecular results

Phylogenetic inference reveals that the species *A. gonogramma* constitutes a distinct lineage separate from the species *A. insons*. The three species delimitation methods identified this species but also identified a subcluster separating the sample BLDNA141. This specimen originates from Colombia while all the other specimens come from Costa Rica. The observed genetic divergence is certainly related to a geographical divergence. A genetic study including samples from more distant localities such as Brazil would better characterize the genetic variability of this species.

**Figures 21-23.**
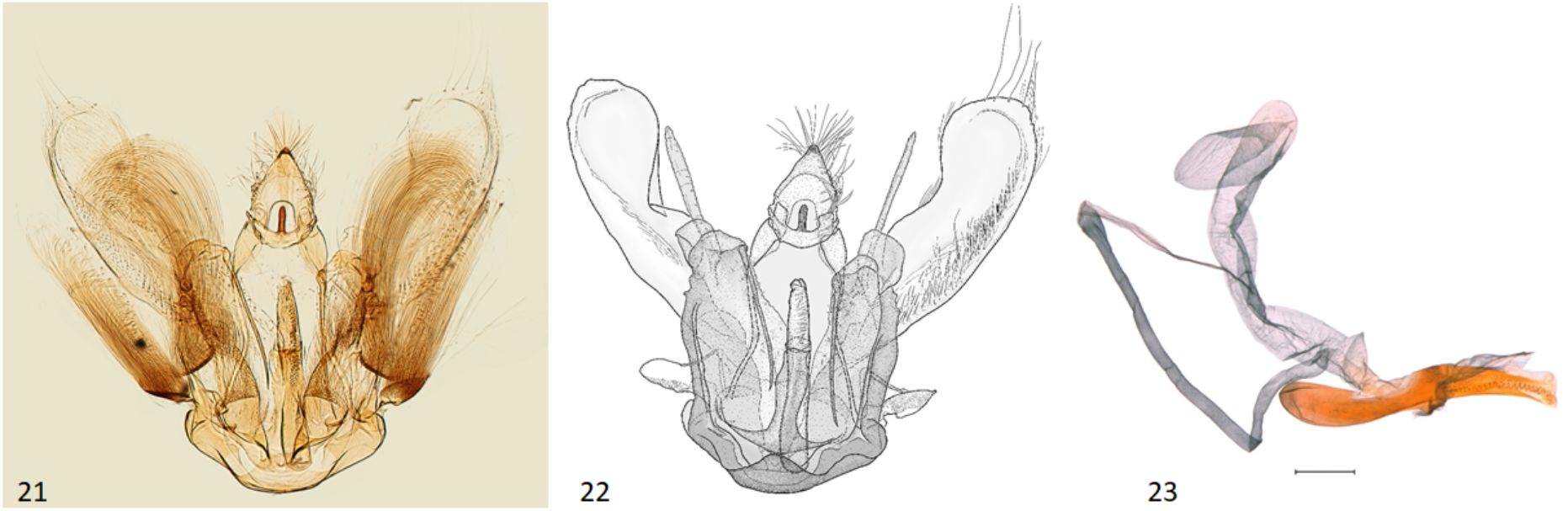
Male genitalia of *Argyria centrifugens* (NMNH) **21** Holotype **22** Drawing of same without pheromone scales **23** Phallus in lateral view (Nicaragua, Selva Negra Ecolodge, MHNG-ENTO-13299); scale bar: 250 µm.

#### Distribution

Fig. 35. Central and South America, from Honduras to Colombia and Brazil. Records from the central west coast of Florida are possibly recent introductions.

#### Remarks

The species was described from a specimen labelled “Type” of an unspecified sex and two other specimens, “Also two others, Cabima, May, 1911 (Busck).” (Dyar 1914). This “Type” is here considered the holotype. There is some variation observed in the male genitalia, especially noticeably in the length of the median process of the gnathos and in the length of the thin pointed rod at the base of the valva.

One female specimen identifiable as *A. centrifugens* was collected in Florida (Largo, Pinellas County), 1 Feb. 1995, by J.G. Filiatrault, deposited in the FSCA (MGCL #1112910). It differs from typical specimens in that the labial palpi are mostly yellowish brown with a few dark brown scales on the first and second palpomeres. However, the maculation is otherwise typical, and the genitalia have the same rugose circumostial chamber as described above.

**Figures 24-26.**
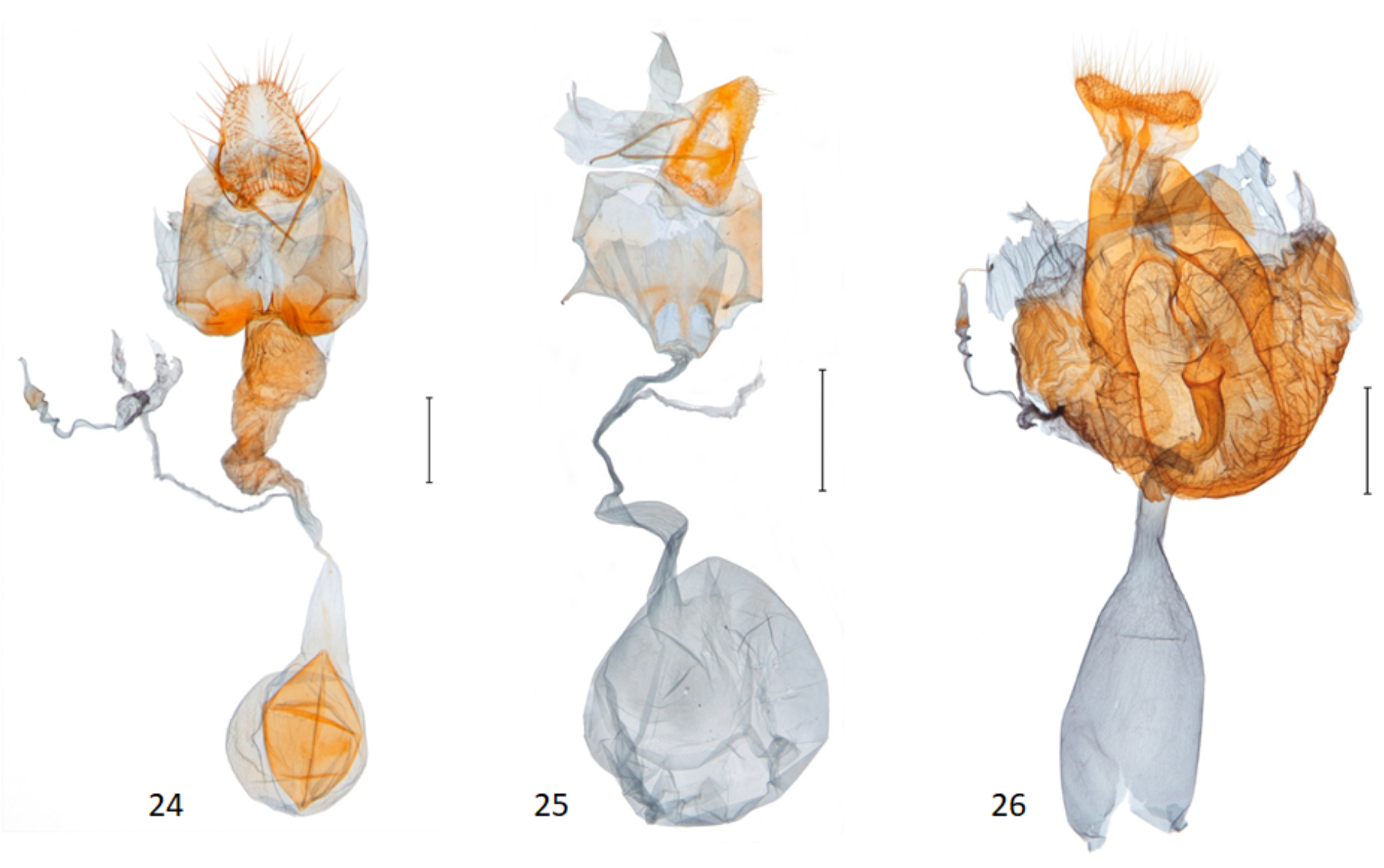
Female genitalia of *Argyria* species **24** *Argyria lacteella* (holotype of *A. abronalis*, with spermatophore inside corpus bursae; OUMNH) **25** *A. diplomochalis* (Anguilla Island, BL 1889, CMNH) **26** *A. centrifugens* (Colombia, Amazonas, Leticia, MHNG-ENTO-97427). Scale bars: 500 µm, except 24, 250 µm.

**Figures 27-28.**
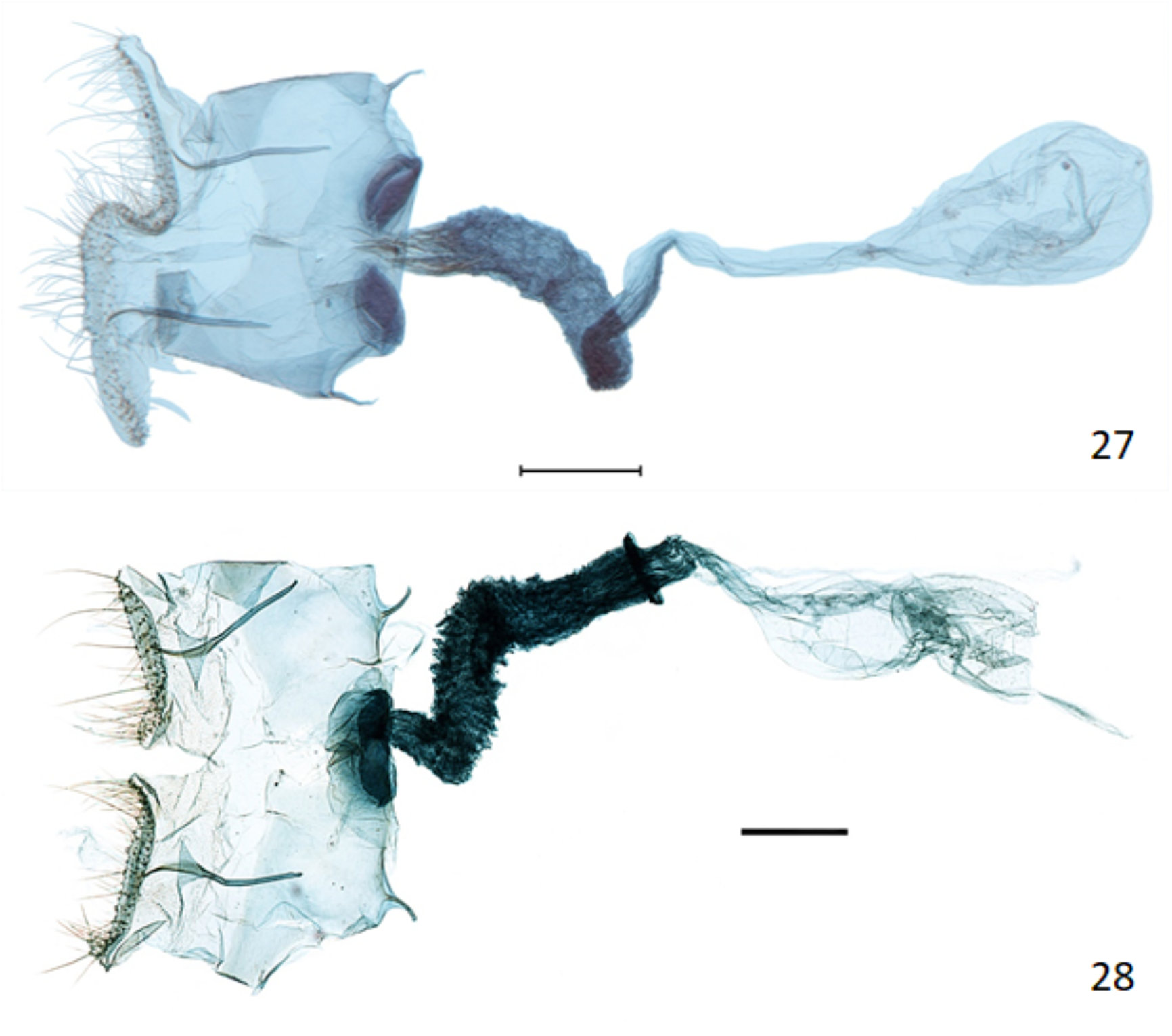
Female genitalia of *Argyria* species with ultimate segments cut dorsomedially from base to apex **27** *Argyria lacteella* (USA, Florida, Glades Co.) **28** *A. gonogramma* (USA, Florida, Orange Co.). Both in FSCA; scale bars: 250 µm.

**Figures 29-32.**
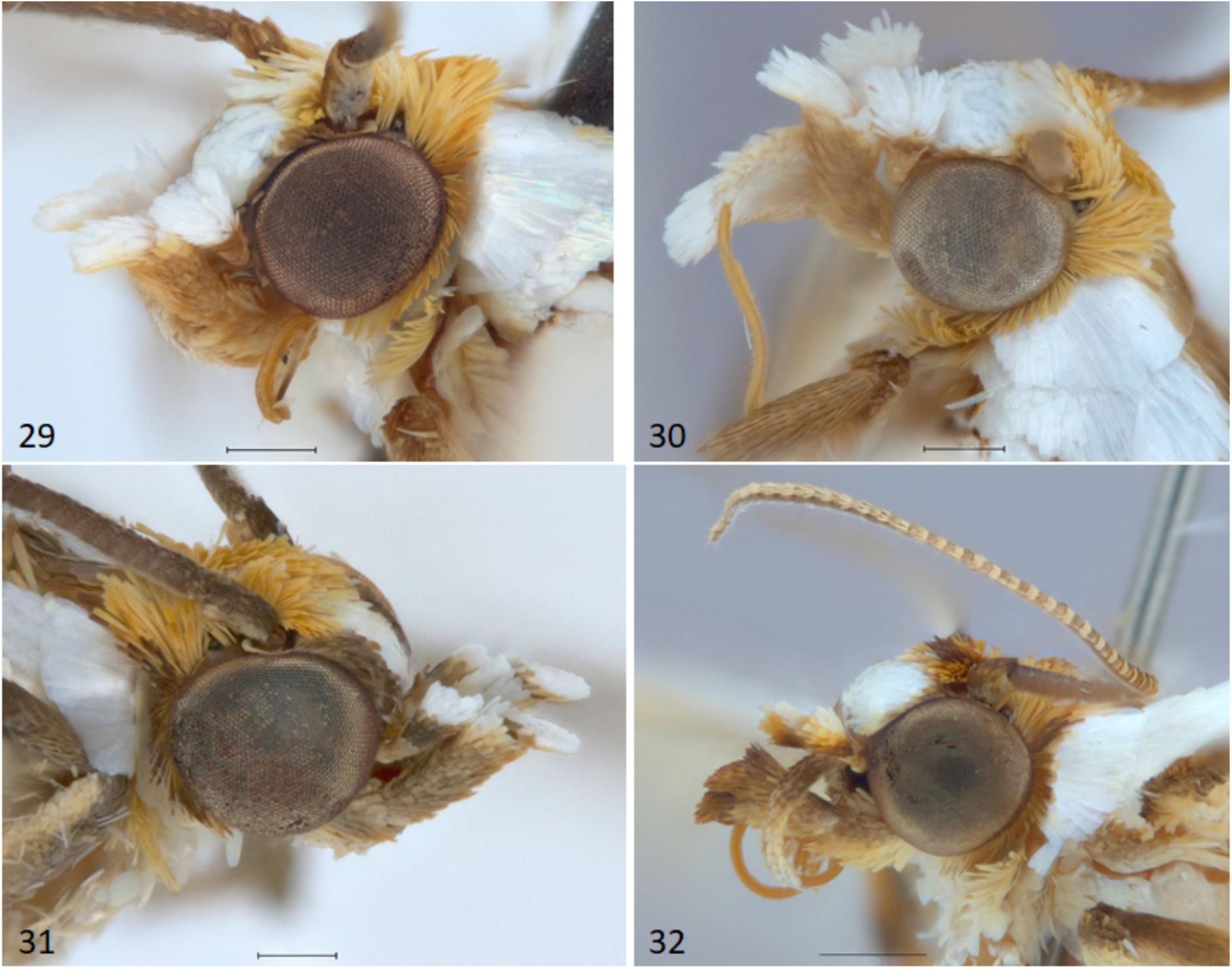
Heads of *Argyria* specimens **29** *A. lacteella* (USA, Florida, Pinellas Co.) **30** *A. gonogramma* (USA, Florida, Levy Co.) **31** *A. diplomochalis* (US Virgin Islands, St. Croix) **32** *A. centrifugens* (Colombia, Amazonas, Leticia). All in MHNG; scale bars: 250 µm except 32, 500 µm.

**Figure 33.**
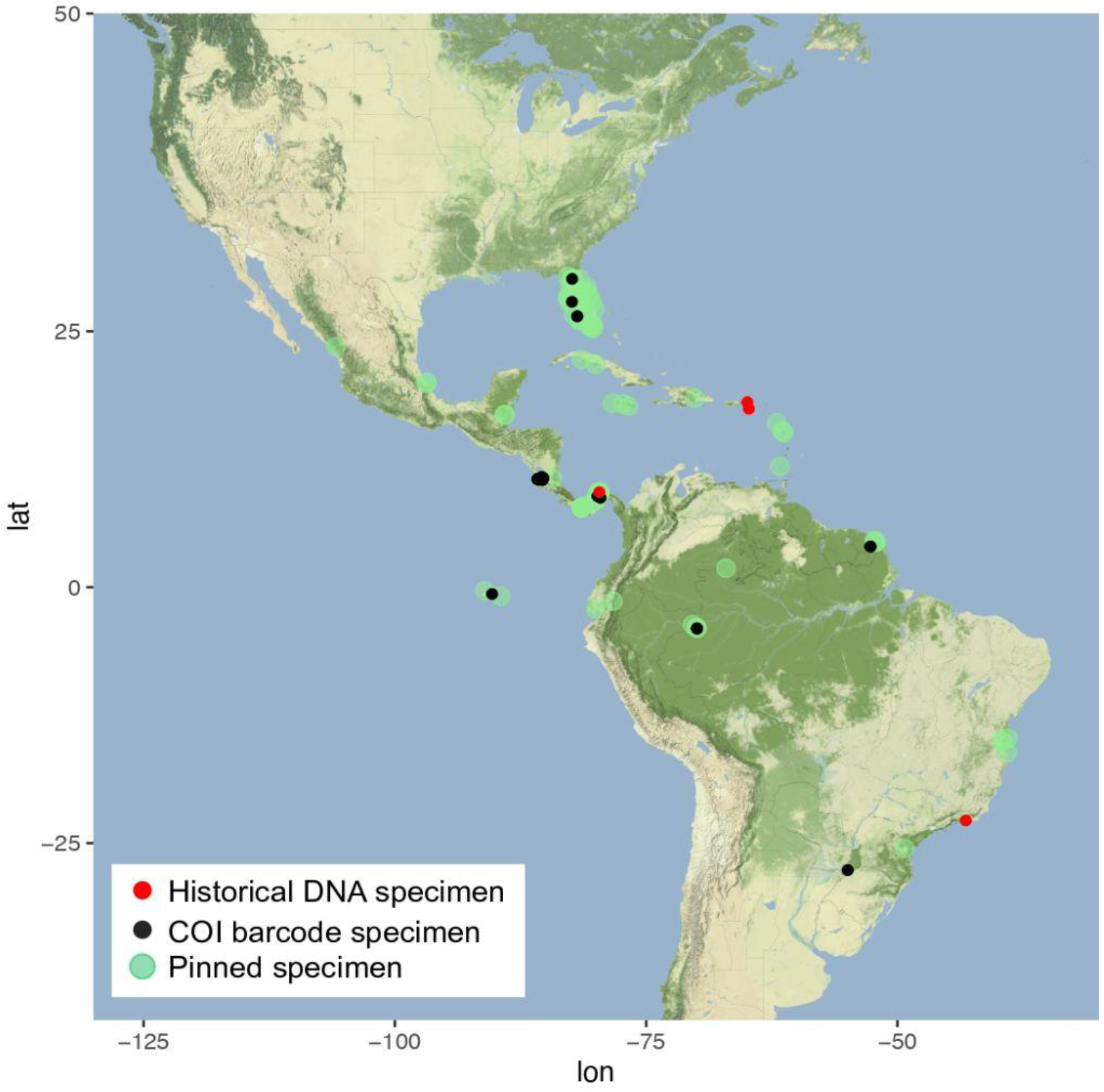
Distribution of *Argyria lacteella* (Fabricius).

**Figure 34.**
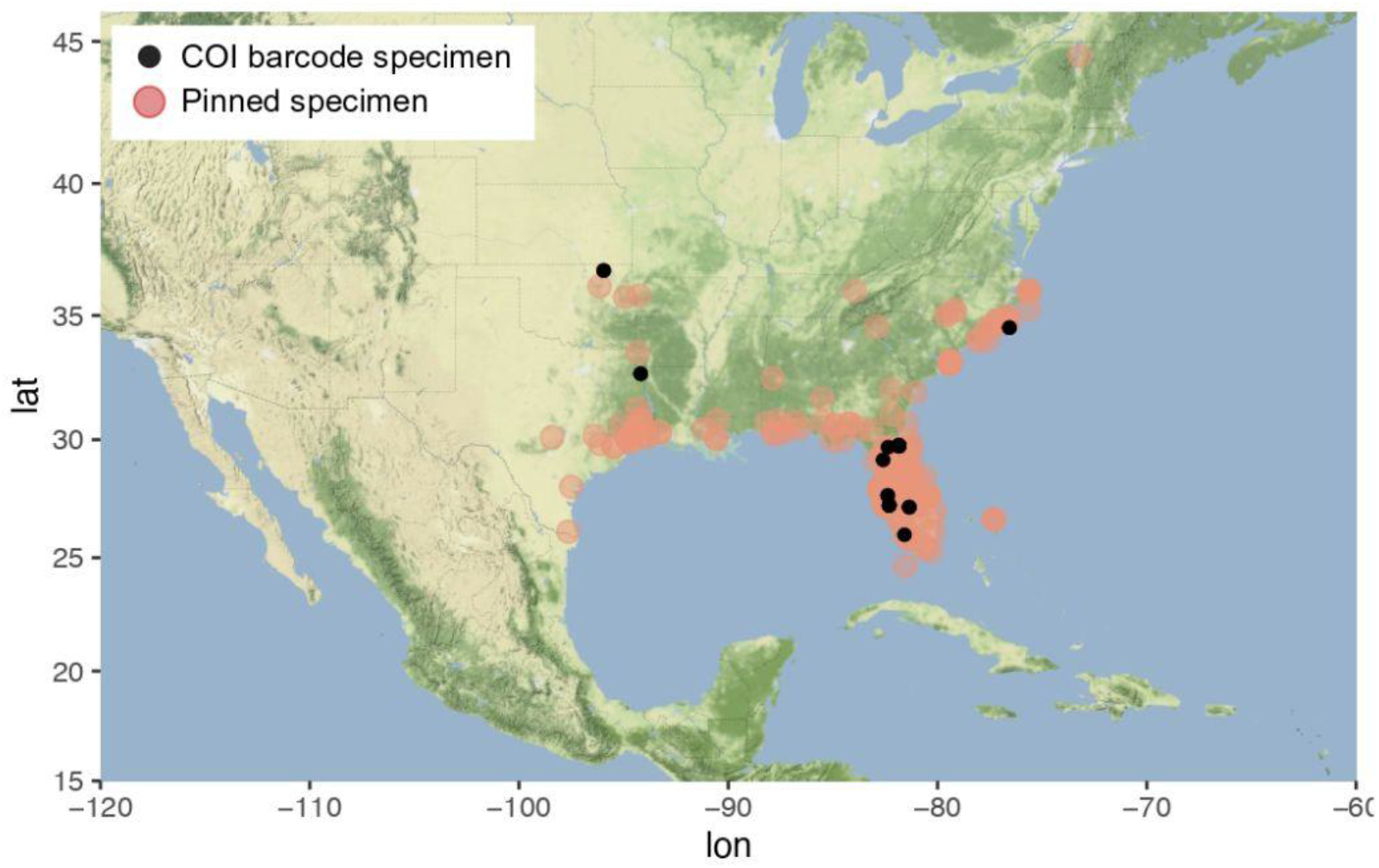
Distribution of *Argyria gonogramma* Dyar.

**Figure 35.**
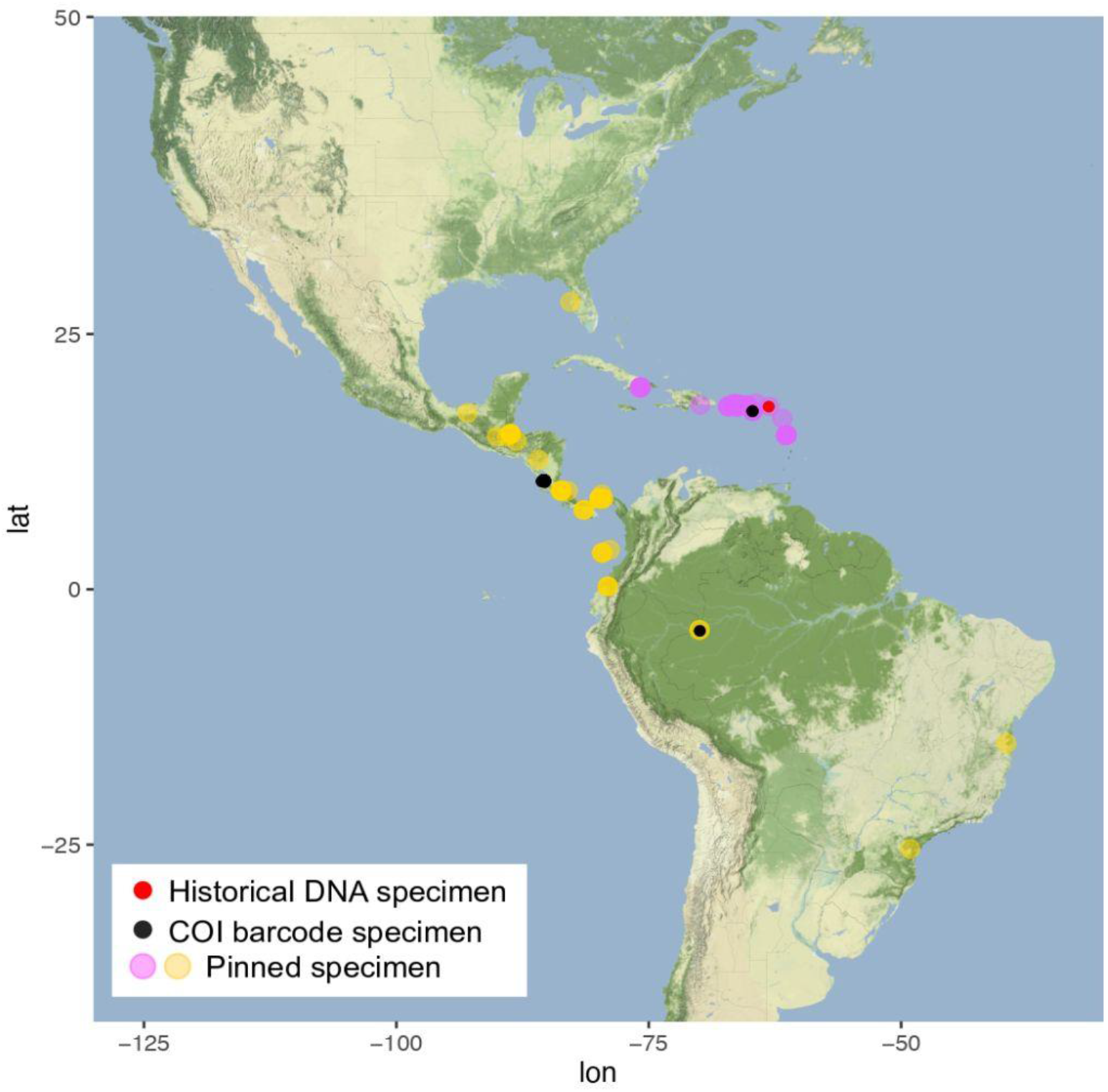
Distributions of *A. centrifugens* Dyar (in yellow) and *A. diplomochalis* Dyar (in pink).

#### General discussion

We were able to resolve complex taxonomic questions for *Argyria* using an innovative DNA hybridization capture protocol to recover high percentages of the DNA barcode of 18^th^- 20^th^ century type specimens. Thus, we were able to solve taxonomic problems regarding synonymies of multiple names applied to the same species. Then we were able to build distribution maps based on refined identities and specimens from multiple museums, leading to other questions regarding responses to environmental change through time. For example, we provided evidence to refine the type locality of *Argyria lacteella* as St Croix Island, whereas the three recent (2021) specimens we examined from that island belong to *A. diplomochalis* (see supplementary material). The possible absence of *A. lacteella* on St Croix currently would not necessarily reflect the environmental situation of 235 years or so ago when the holotype was collected. Various reasons could explain why the latter species was recently collected on St Croix, instead of *A. lacteella*, including habitat change and/or destruction. A thorough moth collecting effort on the island may resolve the question of whether *A. lacteella* still occurs there. Much remains to learn about this group of *Argyria* moths, especially about their biology and immature stages. Their distribution is also incompletely known and some specimen records, for example those of *A. gonogramma* in Vermont and of *A. centrifugens* in Florida need further validation. Finally, many more taxonomic situations such as those in this paper occur in other insect groups that could be resolved using the innovative DNA hybridization capture protocol presented here.

## Acknowledgments

James Hogan (OUMNH) and Thomas Pape (ZMUC) provided two of the type specimens or parts thereof for sequencing and also provided images of type specimens. John Rawlins and James W. Fetzner Jr. (CMNH) provided on loan several hundred specimens of *Argyria* for identification. Ben Proshek (SEL, USDA) provided images of the NMNH type specimens and Floyd Shockley (Collection Manager, Entomology, NMNH) approved the loan of *A. multifacta* to B. Landry during the 2020-2022 covid pandemic that was critical to this study. Carlos Lopez-Vaamonde (INRAE Orléans & IRBI Tours, France) provided sequenced specimens on loan. Paul Hebert and colleagues (Centre for Biodiversity Genomics at the University of Guelph, Ontario, Canada) provided access to BOLD data. John W. Brown (Smithsonian Research Associate, NMNH) sent a gift of two specimens of *Argyria diplomochalis* from St. Croix Island to BL along with several other specimens of *Argyria* from the USA. Ole Karsholt (ZMUC) and Thomas Pape provided information on Fabricius and his work and collection. Steve Nanz provided information on North American species of *Argyria*, and his inquiry about *A. “centrifugens”* in Florida motivated in part the present study. Christina Lehmann-Graber (MHNG) inked and improved the drawing originally made by BL of the holotype of *A. centrifugens*; she also improved the images taken by BL at the MHNG. In the MFNB Robert Schreiber performed the lab work. Sabine Gaal, Lepidoptera Curator (Naturhistorisches Museum Wien, Vienna, Austria), searched for a potential type specimen of *Argyria pusillalis* Hübner. Charles V. Covell, Jr. sorted *Argyria* specimens in the FSCA and MGCL; collecting work in the Bahamas was supported in part by National Geographic Scientific Research Grant # 9439-14, with Principal Investigator Jacqueline Y. Miller. Jean-François Landry (curator of microlepidoptera) and Lisa Bartels opened the Canadian National Collection of Insects, Arachnids, and Nematodes (Ottawa, Ontario) to BL and sent photographs of *Argyria* moths. Steven C. Passoa (USDA/APHIS/PPQ, U.S. Forest Service Northern Research Station and The Ohio State University, Columbus, Ohio, U.S.A.) provided crucial information. We thank them all deeply for their interest and kind support.

## Data availability

Sequence dataset is available on BOLD (DS-ARGYRIA). Raw reads are available on the NCBI SRA XXXXXX (submission upon manuscript acceptance).

## Supplementary Material to

Identity of *Argyria lacteella* (Fabricius) revised

## 1. DNA extractions of historical samples

The protocol is based on Patzold, Zilli and Hundsdoerfer (2020).

### Non destructive lysis

- Remove a leg with forceps and place it in a 1.5 mL tube. Sterilize the forceps on a Steri-250 (Keller; art.n°31100) before leg removal and between each specimen to prevent cross-contamination.
- Add 180 μL of Monarch gDNA Tissue Lysis Buffer and 20 μL of proteinase K. Mix the sample by gently inverting the tube several times and briefly centrifuge to collect droplets. Incubate overnight at 56°C in a Thermomixer at 300 rpm with a heated lid (e.g. ThermoMixerC, Eppendorf).
- Briefly centrifuge to collect droplets and gently pipet out the lysate and transfer it into a 1.5 mL tube for DNA purification.

*N*.*B*.: *The leg (abdomen, or other part of the specimen) can be rinsed with EtOH and dried after the lysis and minimal damage can be observed*.

### Destructive lysis

- Remove a leg with forceps and place it in a 2 mL tube containing Stainless steel beads of 5 mm in diameter (Qiagen/art.n°69989). Sterilize the forceps on a Steri-250 before leg removal and between each specimen to prevent cross-contamination.
- Grind samples 2 times for 2 min at 24 Hz with a TissueLyser (Qiagen). Add 180 μL of Monarch gDNA Tissue Lysis Buffer and 20 μL of proteinase K. Mix the sample by gently inverting the tube several times and briefly centrifuge to collect droplets. Incubate overnight at 56°C in an incubator with a heated lid.
- Centrifuge the lysate at 16’000 x g for 1 min and gently pipet out the lysate and transfer it into a 1.5 mL tube for DNA purification.

### DNA purification

- Add 2 volumes of DNA Cleanup Binding Buffer and mix well by pipetting up and down or flicking the tube, but do not vortex. Briefly centrifuge to collect droplets.
- Add 1.3 volumes of ≥96% Ethanol. Mix well by pipetting up and down and transfer 700 μL of the mix onto the column. Centrifuge for 3 min at 1’000 x g to bind the DNA to the membrane and discard the flow-through. Transfer the remaining mix into the column and centrifuge for 3 min at 1’000 x g. Perform a final centrifugation for 1 min at 16’000 x g and discard the flow-through.
- Add 500 μL of DNA Wash Buffer and spin for 1 min at 16’000 x g. Discard the flow-through.
- Place the column back into the collection tube and repeat this step.
- Place the Monarch DNA Cleanup column back into the collection tube and centrifuge for 2.5 min at 16’000 x g to dry the matrix.
- Transfer the Monarch DNA Cleanup column to a 1.5 ml low-bind tube. Pipet 13 μL of EB Buffer to the center of the matrix. Incubate for 10 min at room temperature. Centrifuge for 1 min at 16’000 x g.
- Pipet the eluate back onto the Monarch DNA Cleanup matrix and repeat the incubation and centrifugation steps.

Quantify the samples with the Qubit™ 1X dsDNA HS assay kit and assess the quality of the DNA with Fragment Analyzer (or Tapestation, or Bioanalyzer).

If necessary, dilute the samples to a maximum of 25-30 ng/μL and proceed with the shotgun libraries preparation protocol.

The concentration of purified DNA extracted from the leg(s) samples, either destructively or not, were estimated based on their fragment analyzer (FA) profiles or measured with a Qubit™ 1X dsDNA HS assay. As they were mainly ranging from ∼0.1-3 ng/μL, no dilution was performed before proceeding to the shotgun library preparation. For CRA01 (DNA extracted with the non-destructive protocol from the abdomen), the concentration of DNA obtained was very high (53.6 ng/µL) and DNA was diluted to ∼27 ng/μL before preparing the shotgun library. The DNA fragment sizes ranged from ∼1 to ≤ 100 ng/μL, with a peak at ∼40 bp. Sample CRA04, the most recently collected, showed a smear from 1 to 800 bp on FA, with a peak at ∼100 bp. The DNA extracted from this leg also showed the highest yield (see Appendix 1 for details).

## 2 Shotgun libraries preparation

The protocol is based on Suchan et al. (2016) with modifications detailed in the supplementary material of Toussaint et al. (2021). The reactions are briefly described here and the modifications compared to Toussaint et al. (2021) outlined in green. In summary, the modifications are the following: the first two purification steps were done with a Monarch DNA & PCR cleanup kit rather than with SPRI beads (ratio beads:sample of 2.7:1) to retain all the small DNA fragments ; for the same reason, the rest of the purification steps were performed with SPRI beads with higher ratios of beads:samples compared to the ratios used in Toussaint *et al*. (2021).

*N*.*B*.: *Unique dual-indexed (UDI) libraries were produced: different and unique P1-barcoded adapters and ILLPCR2 indexing primers were used for each sample (see Appendix 4 for details)*.

*N*.*B*.: *Prepare all solutions (mixes) (and assemble all reactions) at room temperature if not stated otherwise. Samples should be stored at 4°C after each reaction, or at -20°C for long-term storage. Mix solutions (mixes) by pipetting up and down and flicking the tubes or shaking, but never vortex, unless stated otherwise*.

### 1a) Phosphorylation

Take 8 μL of DNA and add a mix of 1 μL of T4 polynucleotide kinase (10 U/μL) and 1 μL of T4 DNA Ligase Buffer (10X) for a total volume of 10 μL. Mix by flicking the tube and briefly centrifuge. Incubate for 30 min at 37°C, 20 min at 65°C.

### 1b) Purification post-phosphorylation with Monarch DNA & PCR cleanup kit (5 μg)

- Add 50 μL of EB buffer to the sample to reach a starting volume of 60 μL.
- Add 2 volumes of DNA Cleanup Binding Buffer (=120 μL) to each tube and transfer into a 1.5 mL tube.
- Mix well by pipetting up and down or flicking the tube. Do not vortex.
- Add 2 volumes of 100% Ethanol (=360 μL), mix by pipetting and transfer to the column.
- Centrifuge for 1 min at 16’000 x g. Discard the flow-through.
- Place the column into the collection tube. Add 200 μL of DNA Wash Buffer and centrifuge for 1 min at 16’000 x g. Discard the flow-through.
- Repeat the Wash Buffer step.
- Place the Monarch DNA Cleanup column back into the collection tube, and centrifuge for 2 min at 16’000 x g to dry the matrix.
- Transfer the Monarch DNA Cleanup column to a 1.5 mL and pipet 12 μL of EB Buffer into the center of the matrix. Incubate for 5 min at room temperature. Centrifuge for 1 min at 16’000 x g (or use gradual centrifugation with a final 1 min centrifugation step at 16’000 x g).
- Pipet the eluate back in the center of the matrix and repeat incubation and centrifugation steps.
- Transfer the eluted sample to a 0.2 mL tube.

#### 2) Heat denaturation

Denaturate the sample at 95°C for 5 min then chill on ice to keep DNA in single strand conformation. Prepare a mix of water + ice blocks and place the 0.2 mL tube in it quickly after taking it out of 95°C.

#### 3) Guanidine tailing

Prepare a mix with 4.7 μL of water, 2 μL of NEBbuffer 4 (10X), 2 μL of CoCl_2_ (2.5 mM), 0.8 μL of GTP (100 mM) and 0.5 μL of Terminal Transferase (TdT) (20 U/μL). Add the mix to the 10 μL of heat-denaturated sample for a total volume of 20 μL. Incubate for 30 min at 37°C and 10 min at 70°C.

#### 4) 2nd strand DNA synthesis

Prepare a mix with 5.4 μL of water, 1 μL of NEBbuffer 4 (10X), 0.6 μL of dNTPs (25 mM each), 1 μL of P2-CCCC oligonucleotide (15 μM) and 2 μL of Klenow Fragment (3’→5’ exo-) (5 U/μL). Add the mix to the 20 μL of guanidine-tailed sample for a total volume of 30 μL. Incubate for 3 hours at 23°C and 20 min at 75°C.

### 5a) Blunt-end reaction

On ice, prepare a mix with 3.95 μL of water, 0.5 μL of NEBbuffer 4 (10X), 0.35 μL of BSA (10 mg/mL) and 0.2 μL of T4 DNA polymerase (3 U/μL). Still on ice, add the mix to the 30 μL of blunt-ended sample for a total volume of 35 μL. Incubate for 15 min at 12°C. Store at 4°C. Only bring to room temperature shortly before the purification step.

### 5b) Purification post-blunt-end reaction with Monarch DNA & PCR cleanup kit (5 μg)

Follow the same protocol than for the purification post-phosphorylation (point 1b), except that only 25 μL of EB buffer is added in the first step to reach a starting volume of 60 μL.

### 6a) Ligation of P1-barcoded adapters

Prepare a mix with 6 μL of water, 2 μL of T4 DNA Ligase Buffer (10X) and 1 μL of T4 DNA Ligase (400 U/μL). Add the mix to the 10 μL of purified post-blunt-end reaction sample. Add 1 μL of P1-barcoded adapters solution (25 μM) for a total volume of 20 μL. Incubate overnight at 16°C.

### 6b) Purification post-ligation reaction with SPRI beads

Purify the sample with SPRI beads according to the producer’s protocol with a beads:sample ratio of 1.4:1. Elute DNA in 20 μL of EB buffer.

### 7a) Adapter fill-in

Prepare a mix with 14.1 μL of water, 4 μL of ThermoPol reaction buffer (10X), 0.4 μL of dNTPs (25 mM each) and 1.5 μL *Bst* DNA polymerase, large fragment (8 U/μL). Add the mix to the 20 μL of purified post-ligation sample for a total volume of 40 μL. Incubate for 20 min at 37°C.

### 7b) Purification post-ligation reaction with SPRI beads

Purify the sample with SPRI beads according to the producer’s protocol with a beads:sample ratio of 2:1. Elute DNA in 20 μL of EB buffer.

### 8a) Indexing PCR

Prepare a mix with 14.1 μL of water, 10 μL of Phusion HF Buffer (5X), 0.4 μL of dNTPs (25 mM each) and 0.5 μL of Phusion U Hot Start DNA Polymerase (2 U/μL). Add 25 μL of the mix to the 20 μL of purified post-adapter fill-in sample. Add 5 μL of ILLPCR2 indexing primer solution (5 μM) for a total volume of 50 μL and mix well. Divide it into two 0.2 mL tubes of 25 μL each. Perform the following 2-steps PCR protocol on a thermocycler with heated lid at 99°C (*ramping = 4°C/s*): incubate at 98°C for 30 s; 30 cycles of 98°C for 7 s and 72°C for 25 s; final elongation for 7 min at 72°C. Pool the 2 PCR reactions together for a total volume of ∼50 μL.

### 8b) Purification of post-PCR reaction with SPRI beads

Purify the sample with SPRI beads according to the producer’s protocol with a beads:sample ratio of 1:1. Elute DNA in 20 μL of water.

POST-PCR DNA concentrations were measured with the Qubit™ 1X dsDNA HS assay kit and ranged from ∼4-16 ng/ μL (see Appendix 2 for details). A drawing of the structure of shotgun libraries can be found in Appendix 3.

## 3. Probe synthesis

Final probe sets were ordered from Integrated DNA Technologies (IDT) using the “50 pmol o-Pool Oligos” option. Both sets consisted of 21 different 133 bases-long oligonucleotides: 108 bases of reference sequence + 25 bases of T7 promoter sequence. The two sets of biotinylated RNA probes synthetized are referred to as “forward probe set” and “reverse probe set”.

Anneal each DNA oligonucleotide probe set to the T7 promoter reverse-complement oligonucleotide in a 20 μL reaction containing annealing buffer, 0.5 μM of each of the 21 oligonucleotide and 10.5 μM of the T7 promoter reverse-complement oligonucleotide (oligonucleotide stock solutions were previously resuspended in EB buffer). Heat each reaction at 95°C for 5 min and anneal into dsDNA by decreasing to 20°C with a down-ramping of 0.1°C/s on a thermocycler with heated lid.

Transcribe into biotinylated RNA probes following HiScribe T7 High Yield RNA Synthesis Kit protocol for “RNA synthesis with Modified Nucleotides” a total of 1 μg of annealed oligonucleotide for each probe set. Use Biotin-16-UTP (10 mM) as the Modified UTP and incubate the reaction for 2 hours at 37°C. Add 1 μL of TURBO™ DNase (2 U/μL) to the RNA probes and incubate 15 min at 37°C. Purify the RNA with RNeasy Mini Kit following a modified version of the “RNA cleanup” protocol: add 675 μL instead of 250 μL of 100% ethanol to the diluted RNA before transferring to the purification column to increase the recovery of small RNA fragments; elute the RNA in 20 μL of RNase-free water; add 1 μL of SUPERase-In™ RNase Inhibitor (20 U/μL) to the purified RNA probes sets.

The concentrations of both probes sets were measured with the Qubit™ RNA HS Assay Kit.

## 4. Enrichment of shotgun libraries with hybridization/capture

The five shotgun libraries produced were quantified with the Qubit™ 1X dsDNA HS assay kit and pooled equimolarly based on their concentration.

### Hybridization/capture with “forward probe set”

Some of the equimolar pool was concentrated with a speedvac at 40°C on Heat+IR mode and centrifugation at 2500 rpm (IR vacuum centrifugal concentrator, catalog n°: NB-503CIR, N-BIOTEK) to a solution of ∼14.3 ng/μL total concentration. Thus, 100 ng of the “forward probe set” were used to capture 100 ng of the concentrated libraries in two separate reactions at 55°C following the hybridization-capture protocol based on the Arbor Biosciences Custom protocol:

Prepare the hybridization mix by mixing 9 μL of SSPE (20X), 0.5 μL of EDTA (500 mM), 3.5 μL of Denhardt’s solution (50X), 3.5 μL of of SUPERase-In™ RNase Inhibitor (20 U/μL), 0.4 μL of water and 5.1 μL of biotinylated RNA probes. Add 0.5 μL of SDS (10%) for a total volume of 22.5 μL and incubate the mix at 65°C in an incubator with heated lid for 5-10 min until dissolution of SDS. Leave the tube with the hybridization mix at room temperature.

Prepare the blocking mix by mixing 2.5 μL of Human Cot-1 DNA (1 mg/mL), 2.5 μL of Salmon sperm DNA (1 mg/mL) and 0.5 μL (= 1 μg total) of RNA blocking oligos BO.P5/BO.P7 for a total volume of 5.5 μL.

Add 5 μL of blocking mix to 7 μL of concentrated shotgun libraries (=∼100 ng) and incubate for 5 min at 95°C in a thermocycler with heated lid at 99°C, then decrease the temperature of the block to 55°C. Place the hybridization mix on the thermocycler and incubate for 5 min at 55°C. Add 18 μL of the heated hybridization mix to the 12 μL of blocking mix and shotgun libraries for a total volume of 30 μL. Mix and transfer to another thermocycler and incubate at 55°C, with heated lid at 65°C, for 40 hours.

Take 30 μL of Dynabeads™ M-280 Streptavidin, place on the magnetic rack and remove supernatant. Take out of the magnetic rack, wash with 200 μL of TEN buffer and mix by pipetting, place on the magnetic rack and remove the supernatant. Perform a total of three washes with 200 μL of TEN buffer and resuspend in 70 μL of TEN buffer. Incubate at 55°C until use.

*N*.*B*.: *The preparation of the Dynabeads™ M-280 Streptavidin should be done the same day that they are used*.

Prepare the wash buffer in excess to account for evaporation and pipetting errors during the preparation of aliquots. For 1 mL of wash buffer solution, mix 8 μL of SDS 10%, 200 μL of

0.1 X SSC/0.1% SDS solution and complete with water up to 1 mL. Aliquot the wash buffer in 0.2 mL tubes containing 185 μL of water buffer and incubate at 55°C at least 45 minutes before use.

*N*.*B*.: *The preparation of the wash buffer and of the 0*.*1 X SSC/0*.*1% SDS solution should be done the same day that they are used*.

After 40 hours of hybridization, add 70 μL of pre-heated Dynabeads™ M-280 Streptavidin to capture the 30 μL of libraries and probes in a total volume of 100 μL. Vortex, short-spin and incubate for 30 min at 55°C. Vortex and centrifuge every 10 min. Place on the magnetic rack, remove the supernatant, add 180 μL of pre-heated wash buffer, vortex, short-spin, then incubate for 10 min at 55°C. Repeat this washing step four times then resuspend in 30 μL of EB buffer and store at RT before the post-capture amplification.

Prepare the post-capture amplification master mix by mixing 10 μL of IS5/IS6 primer solution (5 μM each oligonucleotide) with 10 μL of water and 50 μL of KAPA HiFi HotStart ReadyMix. Add 70 μL of post-capture amplification master mix to 30 μL of captured libraries (still containing the Dynabeads™ M-280 Streptavidin) for a total volume of 100 μL. Mix and divide in two PCR replicates of 50 μL each. Perform the following 3-steps PCR protocol on a thermocycler with heated lid at 99°C (*ramping = 4°C/s*): incubate at 98°C for 2 min; 15 cycles of 98°C for 20 s, 65°C for 30 s and 72°C for 30 s; final elongation for 5 min at 72°C. Pool the two PCR replicates and purify with SPRI beads according to the producer’s protocol with a beads:sample ratio of 0.8:1. Elute DNA in 15 μL of EB buffer.

The two separate hybridization/capture reactions with the “forward probe set” were pooled and concentrated with SPRI beads to 9 μL (in EB buffer) according to the producer’s protocol with a beads:sample ratio of 0.8:1. The concentration measured with the Qubit™ 1X dsDNA HS assay kit was ∼0.9 ng/μL.

7 μL (= 6.1 ng) of the captured-enriched libraries pool was used to perform a second round of hybridization/capture enrichment at 65°C with 100 ng of the “forward probe set”. The protocol was the same than with the hybridization/capture enrichment at 55°C, except for the hybridization, capture with Dynabeads™ M-280 Streptavidin, and wash steps, which were performed at 65°C instead of 55°C. Twenty PCR cycles instead of 15 were used for the post-capture amplification. Libraries were eluted in 15 μL of EB buffer after purification with SPRI beads. The concentration measured with the Qubit™ 1X dsDNA HS assay kit was ∼90 ng/μL.

### Hybridization/capture with “reverse probe set”

The remaining volume of equimolar pool of shotgun libraries was concentrated with SPRI beads according to the producer’s protocol with a beads:sample ratio of 0.8:1 and eluted in 9 μL of EB buffer to maximize the total ng input of shotgun libraries for the hybridization reaction.

The protocol for hybridization/capture was the same than the one used for the hybridization/capture with the “forward probe set” except for the following details:

only one reaction was performed for the hybridization/capture at 55°C and due to the limited material available, only 24.5 ng of concentrated pool of shotgun libraries was used with 100 ng of biotinylated RNA probes.

The concentration of the 15 μL of captured libraries after hybridization/capture at 55°C and enrichment with 15 PCR cycles was ∼0.7 ng/μL. Thus, 4.9 ng (7 μL) of the captured-enriched libraries pool was used to perform a second round of hybridization/capture enrichment at 65°C with 100 ng of the “reverse probe set”. Only 15 PCR cycles instead of 20 were used to amplify the libraries after the hybridization/capture at 65°C. The concentration measured with the Qubit™ 1X dsDNA HS assay kit was ∼13 ng/μL.

## 5. DNA extractions of fresh samples with QIAamp DNA Micro Kit

DNA from two specimens (CRA05 and CRA06) collected in 2021 were extracted with a QIAamp DNA Micro Kit following the protocol for isolation of genomic DNA from tissues with some minor modifications:

- Remove and grind the leg with the same protocol as described above for historical specimens prior to the addition of 180 μL of Buffer ATL and 20 μL of proteinase K.
- After overnight incubation, centrifuge the tube and transfer the lysate to a new 1.5 mL tube taking care not to pipet any non-digested tissues fragments.
- After addition of ethanol, mix the sample by pipetting up and down and transfer directly to the column.
- Elute the DNA in 25 μL of EB buffer and incubate 5-10 min instead of 1 min.

The concentration measured with a Qubit™ 1X dsDNA HS assay kit was ∼0.8 ng/μL for both samples. The profiles obtained with a High Sensitivity Fragment Analyzer 1-6000 bp kit showed a good quality of the DNA with fragments ≥ 6 kb and no fragments < 700 bp.

For the samples BLDNA065, BLDNA137, BLDNA138 and BLDNA141, DNA was extracted using the Macherey-Nagel DNA extraction kit (Dürren, Germany) at the MFNB.

## 6. PCR amplification of fresh DNA samples for Sanger sequencing

The COI barcode was amplified by PCR with the primers pair used in the study of Landry and Andriollo (2020) referred to as H02198 and COImod. PCR amplification parameters needed to be adjusted in regard to the use of so-called “universal primers” and also to the amplification of small amount of DNA. The two following protocols proved to be successful and the required adjustments of PCR parameters are outlined below. PCR samples were purified with SPRI beads according to the producer’s protocol with a beads:sample ratio of 0.8:1; DNA was eluted in 10 μL of EB buffer.

Final PCR protocol with MyFi™ Mix (used for CRA05):

Assemble a 20 μL PCR reaction containing 10 μL of MyFi™ Mix (2X), 0.6 μL of each primer solution (20 μM), ∼1 ng of DNA and water. Perform the following PCR in a thermocycler with heated lid at 99°C and ramping rate at 2°C/s: 3 min of initial denaturation at 95°C, 40 cycles of 30 s denaturation at 94°C, 45 s of annealing at 50°C, 60 s of elongation at 72°C and final elongation for 10 min at 72°C.

Adjustments of PCR parameters compared to a classic PCR with MyFi™ Mix and to the protocol from Landry and Andriollo (2020):

- Ramping rate reduced to 2°C/s for all steps of the PCR.
- Increase of primer concentration to 0.6 μM each instead of 0.4 μM each.
- Increase of initial denaturation time to 3 min at 95°C.
- Increase the total number of cycles to 40.
- Decrease of denaturation temperature from 95°C to 94°C in PCR cycles.
- Reduction of annealing temperature to 50°C.
- In PCR cycles: denaturation time set to 30 s, annealing time extended to 45 s, extension time extended to 60 s.
- Final extension time extended to 10 min.

Final PCR protocol with Platinum™ SuperFi™ DNA Polymerase (used for CRA06)

Assemble a 10 μL PCR reaction containing 2 μL of SuperFi™ Buffer (5X), 0.8 μL of dNTPs (2.5 mM each), 0.5 μL of each primer solution (20 μM), 2 μL 5X SuperFi™ GC Enhancer, 0.1 μL of Platinum™SuperFi™ DNA Polymerase (2 U/μL),∼1 ng of DNA and water. Perform the following PCR in a thermocycler with heated lid at 99°C and ramping rate at 2°C/s: 1 min of initial denaturation at 98°C, 40 cycles of 10 s denaturation at 95°C, 30 s of annealing at 50°C, 30 s of elongation at 72°C and final elongation for 10 min at 72°C.

Adjustments of PCR parameters compared to a classic PCR Platinum™ SuperFi™ DNA Polymerase

- Ramping rate reduced to 2°C/s for all steps of the PCR.
- Increase of primers concentration to 1 μM each instead of 0.5 μM each.
- Increase of initial denaturation time to 1 min at 98°C.
- Increase total number of cycles to 40.
- Decrease of denaturation temperature from 98°C to 95°C in PCR cycles.
- Annealing temperature reduced to 50°C.
- In PCR cycles: denaturation time set to 10 s, annealing time extended to 30 s, extension time extended to 30 s.
- Final extension time extended to 10 min.

For the samples BLDNA065, BLDNA137, BLDNA138 and BLDNA141, the COI barcode region was amplified using the LCO/Nancy primer combination. For samples where amplification failed, two fragments COI-1a (LCO/K699) and COI-1b (COIf220/Nancy) were amplified subsequently. The PCR-mix is composed of 17.80µl of H20, 0.50µl of forward primer (10µM), 0.50µl of reverse primer (10µM), 0.50µl of dNTPs, 1.00µl of Mg, 2.50µl of buffer and 0.20µl of Taq Polymerase. PCR program is composed of an initial temperature of 95°C during 5min followed by 42 cycles of 30s at 95°C, 40s at 49°C, 50s at 72°C and a final step of 10min at 72°C. Amplification results were checked on a 1% agarose gel. Sequencing was done by Macrogen (The Netherlands) in both directions. Sequences were eye-checked and aligned using Phyde 0.9971 (Müller et al., 2005).

## 8. Acknowledgements

We thank Marjorie Labédan (University of Lausanne) for her help in the development of this protocol, especially in the tests to adapt the shotgun library protocol to very degraded DNA samples extracted with Monarch DNA & PCR cleanup kit. We thank Hélène Mottaz (Muséum d’histoire naturelle, Geneva) for her help in revising this protocol.

## 9. Appendix

## Appendix 1: Information on extracted DNA

| ID | Tissue | Year of collect | Extraction method | DNA conc. in ng/μL (based on) | DNA fragment sizes: range in bp | DNA fragment sizes: peak in bp |
| --- | --- | --- | --- | --- | --- | --- |
| CRA 01 | abdomen | before 1859 | non-destructive | 53.6 (Qubit) | 1 to 100 | 30-35 |
| CRA 02 | leg | before 1863 | non-destructive | 0.4 (FA) | 1 to 100 | 30-35 |
| CRA 03 | leg | 1784-1789 | destructive | 0.5 (FA) | 1 to 100 | 30-35 |
| CRA 04 | leg | 1992 | destructive | 2.94 (FA) | 1 to 800 | 100 |
| CRA 07 | 2 legs | 1911 | destructive | 0.17 (FA) | 1 to 100 | 30-35 |

## Appendix 2: Information on shotgun libraries

| ID | DNA conc. in ng/μL POST-PCR mesured with Qubit |
| --- | --- |
| CRA01 | 15.8 |
| CRA02 | 6.67 |
| CRA03 | 5.91 |
| CRA04 | 4.67 |
| CRA07 | 4.72 |

## Appendix 3: Final POST-PCR product of shotgun library protocol

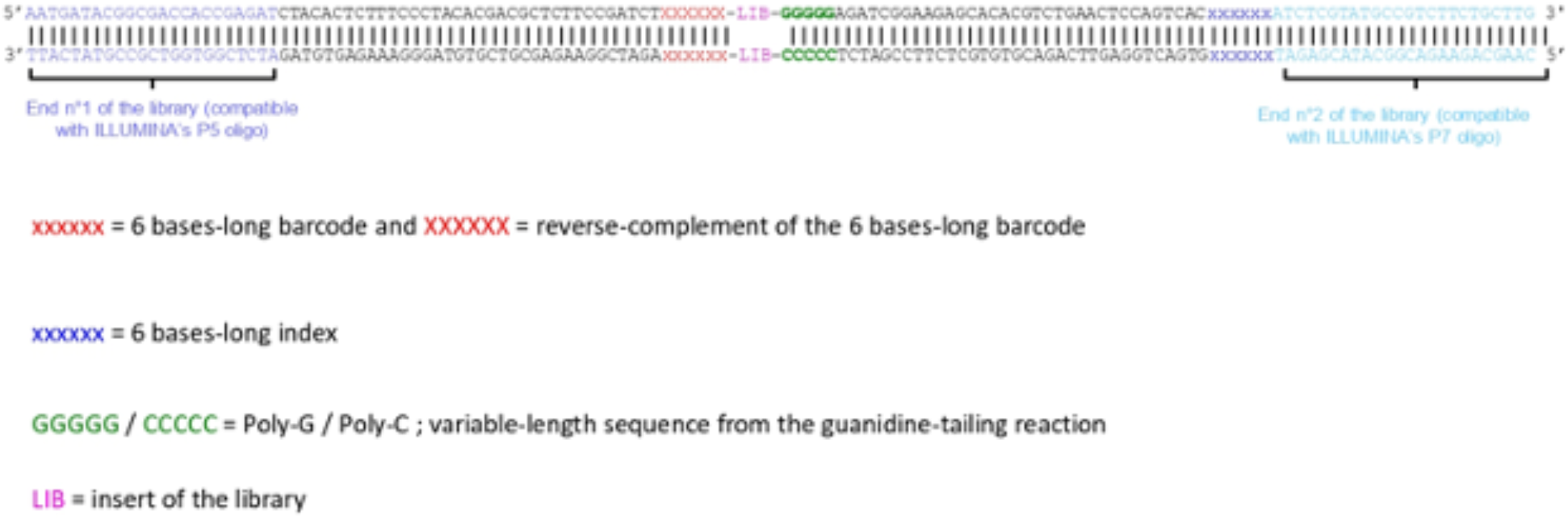

## Appendix 4: Preparation of oligonucleotides solutions

P2-CCCC solution (15 μM):

**Sequence 5’->3’ P2-CCCC oligonucleotide:**

GTGACTGGAGTTCAGACGTGTGCTCTTCCGATCTCCCCC

**Preparation:**

Prepare a 15 μM solution in Tris-HCl 10 mM.

P1-barcoded adapters solution (25 μM)

**Sequence 5’->3’ P1-X.1-barcoded oligonucleotide:**

ACACTCTTTCCCTACACGACGCTCTTCCGATCTXXXXXX

**Sequence 5’->3’ P1-X.2-barcoded oligonucleotide:**

xxxxxxAGATCGGAAGAGC

where xxxxxx = 6 bases-long barcode and XXXXXX = reverse-complement of the 6 bases-long barcode.

**Preparation**

Prepare a 25 μM each oligonucleotide solution in annealing buffer. Heat at 95°C for 5 min and anneal into dsDNA by decreasing to 20°C with a down-ramping of 0.1°C/s on a thermocycler with heated lid. Prepare a different solution using a different barcode sequence for each sample if libraries are designed in UDI.

ILLPCR2 indexing primer solution (5 μM)

**Sequence 5’->3’ ILLPCR1 oligonucleotide:**

AATGATACGGCGACCACCGAGATCTACACTCTTTCCCTACACGACG

**Sequence 5’->3’ ILLPCR2_X oligonucleotide:**

CAAGCAGAAGACGGCATACGAGATxxxxxxGTGACTGGAGTTCAGACGTGTGC

where xxxxxx = 6 bases-long index

**Preparation**

Prepare a 5 μM each oligonucleotide solution in Tris-HCl 10 mM.

RNA blocking oligos BO.P5/BO.P7 (2 μg total/μL)

**Sequence 5’->3’ T7 promoter oligonucleotide:**

AGTACTAATACGACTCACTATAGG

**Sequence 5’->3’ BO.P5 oligonucleotide:** AGATCGGAAGAGCGTCGTGTAGGGAAAGAGTGTAGATCTCGGTGGTCGCCGTATCATTCCCTATAGTGAGTCGTATTAGTACT

**Sequence 5’->3’ BO.P7 oligonucleotide:** AGATCGGAAGAGCACACGTCTGAACTCCAGTCACNNNNNNATCTCGTATGCCGTCTTCTGCTTGCCCTATAGTGAGTCGTATTAGTACT

where NNNNNN = DNA Inosine base

**Preparation**

Prepare a solution of T7 promoter and BO.P5 10 μM each oligonucleotide in annealing buffer. Heat at 95°C for 5 min and anneal into dsDNA by decreasing to 20°C with a down-ramping of 0.1°C/s on a thermocycler with heated lid. Transcribe 0.7 μL of annealed oligonucleotide solution into RNA following HiScribe T7 High Yield RNA Synthesis Kit protocol for “RNA synthesis with Modified Nucleotides” with a 2 hours incubation at 37°C. Add 1 μL of TURBO™ DNase (2 U/μL) to the RNA and incubate 15 min at 37°C. Purify the RNA with RNeasy Mini Kit following a modified version of the “RNA cleanup” protocol: add 675 μL instead of 250 μL of 100% ethanol to the diluted RNA before transferring to the purification column to increase the recovery of small RNA fragments; elute the RNA in 20 μL of RNase-free water; add 1 μL of SUPERase-In™ RNase Inhibitor (20 U/μL) to the purified BO.P5 RNA blocking solution.

Follow the same protocol with T7 promoter and BO.P7 oligonucleotide to obtain a BO.P7 RNA blocking solution.

Quantify both BO.P5 and BO.P7 RNA blocking solutions with a Qubit™ RNA HS assay kit and prepare an equimolar pool of 2 μg total RNA per μL to be used as RNA blocking oligos BO.P5/BO.P7 solution in the hybridization/capture protocol.

IS5/IS6 primer solution (5 μM)

**Sequence 5’->3’ IS5 oligonucleotide:**

AATGATACGGCGACCACCGAGAT

**Sequence 5’->3’ IS6 oligonucleotide:**

CAAGCAGAAGACGGCATACGAGAT

**Preparation:**

Prepare a 5 μM each oligonucleotide solution in Tris-HCl 10 mM.

Sanger sequencing primers (20 μM)

**Sequence 5’->3’ H02198 oligonucleotide:**

TAAACTTCAGGGTGACCAAAAAATCA

**Sequence 5’->3’ COImod oligonucleotide:**

AGTTCTAATCATAARGATATYGG

**Preparation**

Prepare a 20 μM solution in Tris-HCl 10 mM for both Sanger sequencing primers.

## Appendix 5: List of reagents

Appendix 5: List of reagents:
| REAGENT | CATALOG NUMBER | PRODUCER |
| --- | --- | --- |
| Annealing buffer (100 mM Tris-HCl, 500 mM EDTA) | Homemade from stock solutions (Invitrogen) |  |
| Bio-16-UTP (10 mM ) | AM8452 | Invitrogen (Thermo Fisher Scientific) |
| BSA (10 mg/mL) | B9000S=20 mg/ml diluted to 10 mg/ml | New England Biolabs |
| *Bst* DNA Polymerase, Large Fragment (8 U/μL) | M0275S, M0275L | New England Biolabs |
| CleanNGS purification beads | CNGS0500 = 1 bottle 500 mL | CleanNA |
| CoCl<sub>2</sub> (2.5 mM) | B0252S (provided with TdT M0315) | New England Biolabs |
| Denhardt's solution 50X | 750018 | Invitrogen (Thermo Fisher Scientific) |
| dNTP Set, 100 mM Solutions | R0181, R0182, R0186 | Thermo Scientific |
| Dynabeads™ M-280 Streptavidin | 11205D, 11206D | Invitrogen (Thermo Fisher Scientific) |
| EB Buffer (=10 mM Tris-Cl, pH 8.5) | 19086 | QIAGEN |
| EDTA (0.5 M), pH=8 | R1021 | Thermo Scientific |
| EDTA (0.5 M), pH=8, Ultrapure | 15575020 | Invitrogen (Thermo Fisher Scientific) |
| Ethanol absolute anhydrous RPE - For analysis - ACS - Reag. Ph.Eur. - Reag. USP 2.5 L | 4146082 | Carlo Erba Reagents |
| GTP solution, Tris buffered (100 mM) | R1461 | Thermo Scientific |
| High Sensitivity NGS Fragment Analysis Kit (1 - 6000bp) | DNF-474-0500 | Agilent Technologies |
| HiScribe T7 High Yield RNA Synthesis Kit, 50 reactions | E2040S | New England Biolabs |
| Human Cot-1 DNA™ (1 mg/mL) | 15279011 | Invitrogen (Thermo Fisher Scientific) |
| KAPA HiFi HotStart Ready Mix, 1.25 ml | KK2601 (07958927001), KK2602 (07958935001) | Kapa Biosystems (Roche Diagnostics) |
| Klenow Fragment (3'→5' exo-) (5 U/μL) | M0212S, M0212L | New England Biolabs |
| Molecular biology grade water HyClone 1 L | SH30538.03 | Cytiva Europe GmbH |
| Monarch DNA & PCR cleanup kit (5 μg) | T1030 | New England Biolabs |
| Monarch gDNA Tissue Lysis Buffer | T3010 | New England Biolabs |
| MyFi™ Mix | BIO-25049, BIO-25050 | Bioline |
| NaCl (5 M), RNase-free | AM9760G, AM9759 | Invitrogen (Thermo Fisher Scientific) |
| NEBuffer 4 10X | B7004S | New England Biolabs |
| Phusion HF Buffer 5X | F-518 (provided with Phusion U Polymerase F555) | Thermo Scientific |
| Phusion U Hot Start DNA Polymerase (2 U/μL) | F555S, F555L | Thermo Scientific |
| Platinum™ SuperFi™ DNA Polymerase (2U/μL) | 12351010, 12351050, 12351250 | Invitrogen (Thermo Fisher Scientific) |
| Proteinase K (20 mg/mL) | 19131, 19133 | QIAGEN |
| QIAamp DNA Micro Kit (50) | 56304 | QIAGEN |
| Qubit™ 1X dsDNA HS assay Kit | Q33230, Q33231 | Invitrogen (Thermo Fisher Scientific) |
| Qubit™ RNA HS Assay Kit | Q32852, Q32855 | Invitrogen (Thermo Fisher Scientific) |
| RNeasy Mini kit | 74104, 74106 | Qiagen |
| Salmon Sperm DNA (1 mg/mL) | 15632011 | Invitrogen (Thermo Fisher Scientific) |
| SDS (10%), BioUltra, for molecular biology | 71736 | Sigma-Aldrich |
| SPRI beads -> CleanNGS purification beads | CNGS0500 = 1 bottle 500 mL | CleanNA |
| SSC 20X | 15557044, 15557036 | Invitrogen (Thermo Fisher Scientific) |
| SSPE 20X, Ultrapure | 15591043 | Invitrogen (Thermo Fisher Scientific) |
| SUPERase-In™ RNase Inhibitor (20 U/μL) | AM2694, AM2696 | Invitrogen (Thermo Fisher Scientific) |
| SuperFi™ Buffer 5X | 12355005, Provided with Platinum SuperFi DNA Polymerase | Thermo Scientific (Thermo Fisher Scientific) |
| SuperFi™ GC Enhancer 5X | Provided with Platinum SuperFi™ DNA Polymerase | Thermo Scientific (Thermo Fisher Scientific) |
| T4 DNA Ligase (400 U/μL) | M0202S, M0202L | New England Biolabs |
| T4 DNA Ligase Reaction Buffer 10X (10mM ATP) | B0202S (provided with T4 DNA Ligase M0202) | New England Biolabs |
| T4 DNA Polymerase (3 U/μL) | M0203S, M0203L | New England Biolabs |
| T4 Polynucleotide Kinase (10 U/ μL) | M0201S, M0201L | New England Biolabs |
| TEN (10 mM Tris-HCl, 1 mM EDTA, 1 M NaCl) | Homemade from stock solutions (Invitrogen) |  |
| Terminal Transferase (TdT) (20 U/ μL) | M0315S, M0315L | New England Biolabs |
| ThermoPol Reaction Buffer Pack 10X | B9004 (provided with *Bst* DNA Polymerase Large Fragment M0275) | New England Biolabs |
| Tris (1 M), pH 8.0, RNase-free | AM9855G, AM9856 | Invitrogen (Thermo Fisher Scientific) |
| TURBO™ DNase (2 U/μL) | AM2238, AM2239 | Invitrogen (Thermo Fisher Scientific) |

